# Deep learning enables cross-species annotation and attribution of ageing states in haematopoietic stem and immune cells

**DOI:** 10.64898/2026.07.24.730582

**Authors:** Simiao Zhao, Bowen Zhang, Melinda Czéh-Bhardwaj, Xuehao Zhai, Niels Asger Jakobsen, Christopher Yau, Pietro Liò, Claus Nerlov

## Abstract

Mouse single-cell ageing studies provide experimentally controlled age contrasts, but using mouse-labelled data to annotate human ageing states is limited by species, donor and assay effects in sparse transcriptomic and chromatin profiles. We developed a cross-species annotation workflow that treats mouse-to-human prediction as a target-validated domain-adaptation problem. The workflow uses orthologue-aligned features, a residual encoder, an age classifier and a species discriminator trained with two-phase adversarial optimisation, and couples prediction with stability-based gene attribution. In haematopoietic stem cells (HSCs), the model achieved held-out human AUROCs of 0.933 in scRNA-seq and 0.953 in scATAC-seq. In an independent CD8+ T-cell scRNA-seq setting, the held-out human AUROC was 0.941. Ablation analyses indicated that residual connections, ELU activation and two-phase training improved predictive performance and attribution stability. Consensus attributions from DeepLIFT, Integrated Gradients and saliency recovered conserved ageing-associated genes with greater cross-species overlap than differential expression alone. In a COVID-19 convalescent cohort, severe disease in younger adults was associated with a higher fraction of CD8+ cells classified as old-like by the pretrained model. These results support a reproducible framework for testing, interpreting and releasing cross-species single-cell ageing models, while highlighting the need for target-domain validation when mouse labels are transferred to human data.

## Introduction

Ageing is not a single molecular process but a progressive reorganisation of tissue maintenance, inflammatory tone, metabolic state and regenerative capacity (ref. 1). Haematopoietic stem cells (HSCs) sit near the centre of this problem. They sustain lifelong blood production, yet ageing changes their self-renewal, lineage output and clonal behaviour, with recurrent shifts towards myeloid and platelet-biased states (refs. 2–4). Ageing also remodels mature immune compartments. In CD8+ T cells, mouse and human studies have identified age-associated expansion, clonal skewing and altered effector-memory states, including conserved GZMK+ and other inflammatory programmes (refs. 5,6). These compartments are therefore biologically informative, but they are also hard to study directly in humans because relevant cells are rare, sampling is invasive and longitudinal perturbation experiments are usually impossible.

Mouse systems address part of the experimental problem. Age, genotype, environment and tissue collection can be controlled, and single-cell profiling can be performed at scales that are unrealistic in human cohorts. The difficulty is that mouse ageing is not a small, noisy version of human ageing. Species differ in lifespan, immune ecology, marrow composition, cell-state frequencies and regulatory sequence architecture. Even after one-to-one orthologue mapping, a gene-expression or chromatin-accessibility matrix contains both ageing signal and species signal. A model that performs well within mouse data may therefore be using molecular rules that do not transfer to human cells. Conversely, a purely descriptive comparison may identify conserved pathways without testing whether those pathways are sufficient for age-state annotation in held-out human samples.

Single-cell genomics makes this tension visible. Large mouse and human atlases show that ageing effects are cell-type specific rather than uniform across tissues (refs. 6,7). Single-cell RNA sequencing measures transcript abundance at cell resolution, whereas single-cell ATAC-seq and gene-activity scores provide a regulatory view of the same biological problem. These data types are sparse, high-dimensional and strongly affected by donor, platform and batch structure. Integration methods such as canonical-correlation approaches and Harmony have made it possible to align cells across datasets and conditions (refs. 8,9). Alignment alone, however, does not define the supervised transfer problem addressed here. The relevant test is whether age labels learned from mouse cells can annotate independent human cells of the same broad lineage without the model simply learning a species classifier.

Computationally, this is a domain-adaptation problem. The source domain contains labelled mouse cells; the target domain contains human cells whose distribution differs from the source because of species, donor and assay effects. Classical supervised learning assumes that training and test examples are drawn from comparable distributions, an assumption that is often violated in cross-species single-cell analysis (refs. 10,11). Reference-mapping methods in single-cell genomics have shown that transfer learning can be useful when query data must be related to an existing atlas, but they also illustrate the need to preserve biological state rather than remove all variation indiscriminately (ref. 12). In this setting, age is the biological label to be transferred, whereas species is a nuisance domain that can create shortcut solutions if the model exploits features that separate mouse from human more strongly than young from old (ref. 13).

Domain-adversarial neural networks (DANNs) provide one way to formalise this objective (ref. 14). A shared encoder maps each cell into a latent representation. An age-classification head is trained to predict young versus old, while a domain-discrimination head is trained to predict mouse versus human. The encoder is then pressured to retain information that supports age classification while reducing information that allows species discrimination. Biologically, the objective is not to hide meaningful variation from the model, but to discourage solutions driven by species-specific markers rather than conserved age-associated features. Computationally, the procedure defines a constrained representation-learning problem in which label predictiveness and domain invariance are optimised together under distribution shift.

This formulation is not automatically reliable in single-cell data. In single-cell data, age and species can be partially entangled, class sizes can be imbalanced, and adversarial gradients can destabilise training. Over-correction may erase the biology that the model is supposed to transfer. Under-correction leaves species structure in the embedding and inflates apparent performance in source-like settings. Architectural choices therefore matter. Residual connections improve optimisation in deep networks, batch normalisation and dropout regularise high-dimensional models, and ELU activations can reduce saturation during training (refs. 15–18). We used these elements in a residual multilayer perceptron encoder and paired them with a two-phase optimisation scheme that separates age-classifier updates from adversarial domain updates. The ablation analyses below test whether these choices affect both predictive performance and attribution stability, rather than treating them as arbitrary implementation details.

High predictive performance is insufficient for an ageing resource. A classifier can rank samples correctly while relying on features that are hard to interpret or inconsistent across runs. Differential expression provides an important baseline because it identifies genes with marginal age-associated changes within each species, but it does not ask which features the trained cross-species model actually uses. Attribution methods such as DeepLIFT, Integrated Gradients and saliency maps estimate how input genes contribute to a model output, and Captum provides a common implementation framework for these analyses (refs. 19–22). Because attribution from a single trained model can be unstable, we aggregate repeated runs and focus on consensus genes whose importance is reproducible across methods and species. This converts the model from a black-box classifier into a generator of testable conserved-feature hypotheses.

Here we develop and evaluate a cross-species ageing-state annotation workflow for haematopoietic stem and immune cells. The workflow combines orthologue-aligned single-cell features, a residual two-phase DANN, target-domain validation, baseline comparisons, architecture ablations and stability-based attribution. We test the framework in HSC scRNA-seq and HSC scATAC-seq, evaluate transfer in an independent CD8+ T-cell scRNA-seq setting and apply the pretrained CD8+ model to a COVID-19 convalescent cohort as an external immune-state readout. Together, these analyses test whether mouse-labelled single-cell data can support human ageing-state annotation when models are explicitly evaluated for transfer, constrained against species shortcuts and interrogated for reproducible genes.

## Results

### A two-phase framework learns transferable ageing-state embeddings across species

We organised the study around a shared latent representation that receives human and mouse HSC gene-expression or gene-activity matrices and feeds two heads: an age classifier and a domain discriminator (Fig. 1a,b). The encoder uses residual fully connected blocks with batch normalisation, dropout and ELU activations, so that age-discriminative signal is learned in a compact embedding rather than directly from raw high-dimensional features (refs. 15–18). Instead of the usual gradient-reversal update, optimisation alternates between two steps (Fig. 1c): one phase updates the encoder and age head on labelled mouse cells, whereas the second phase updates the domain head while adversarially adjusting the encoder to reduce species information.

**Figure 1.**
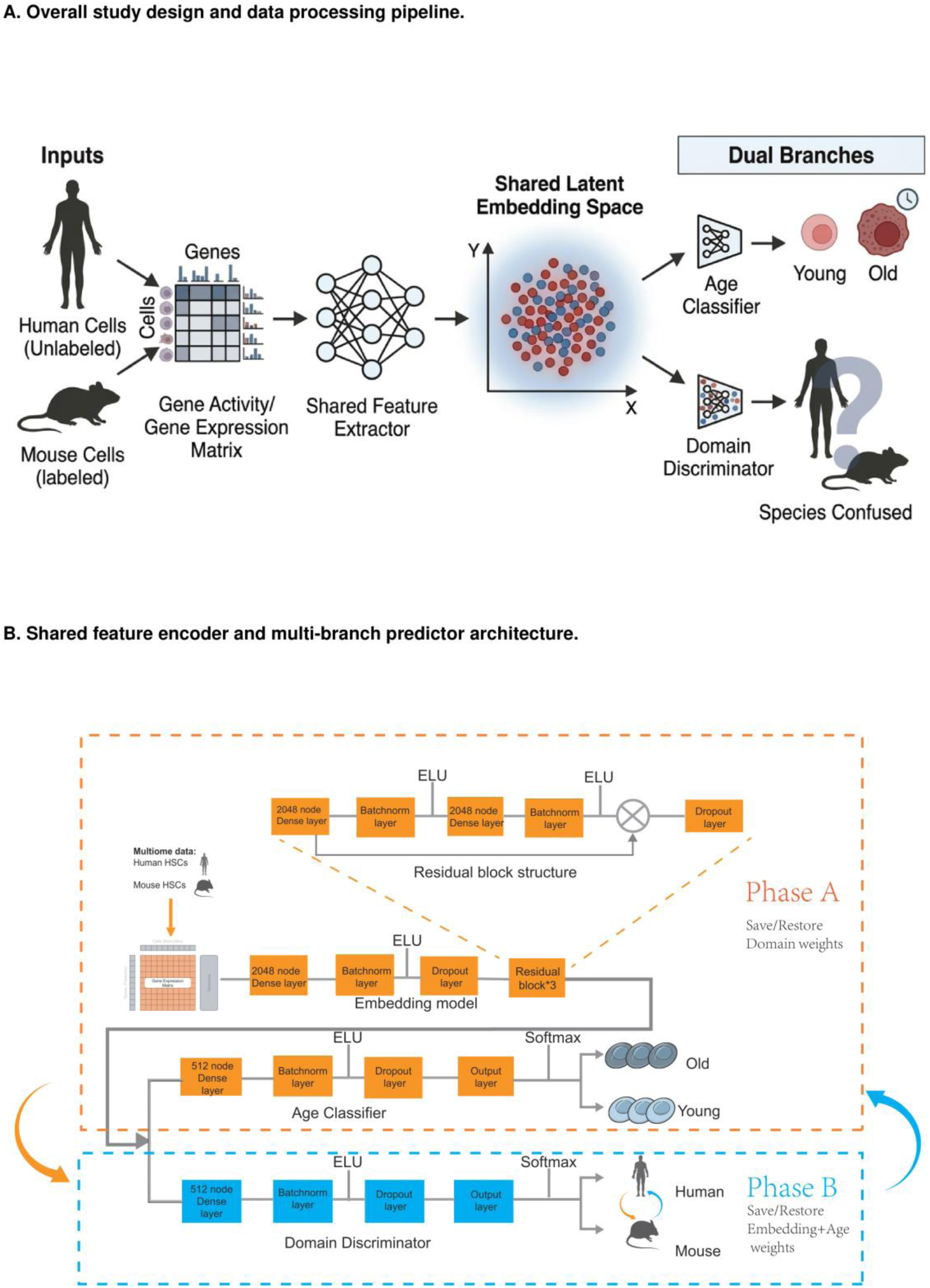
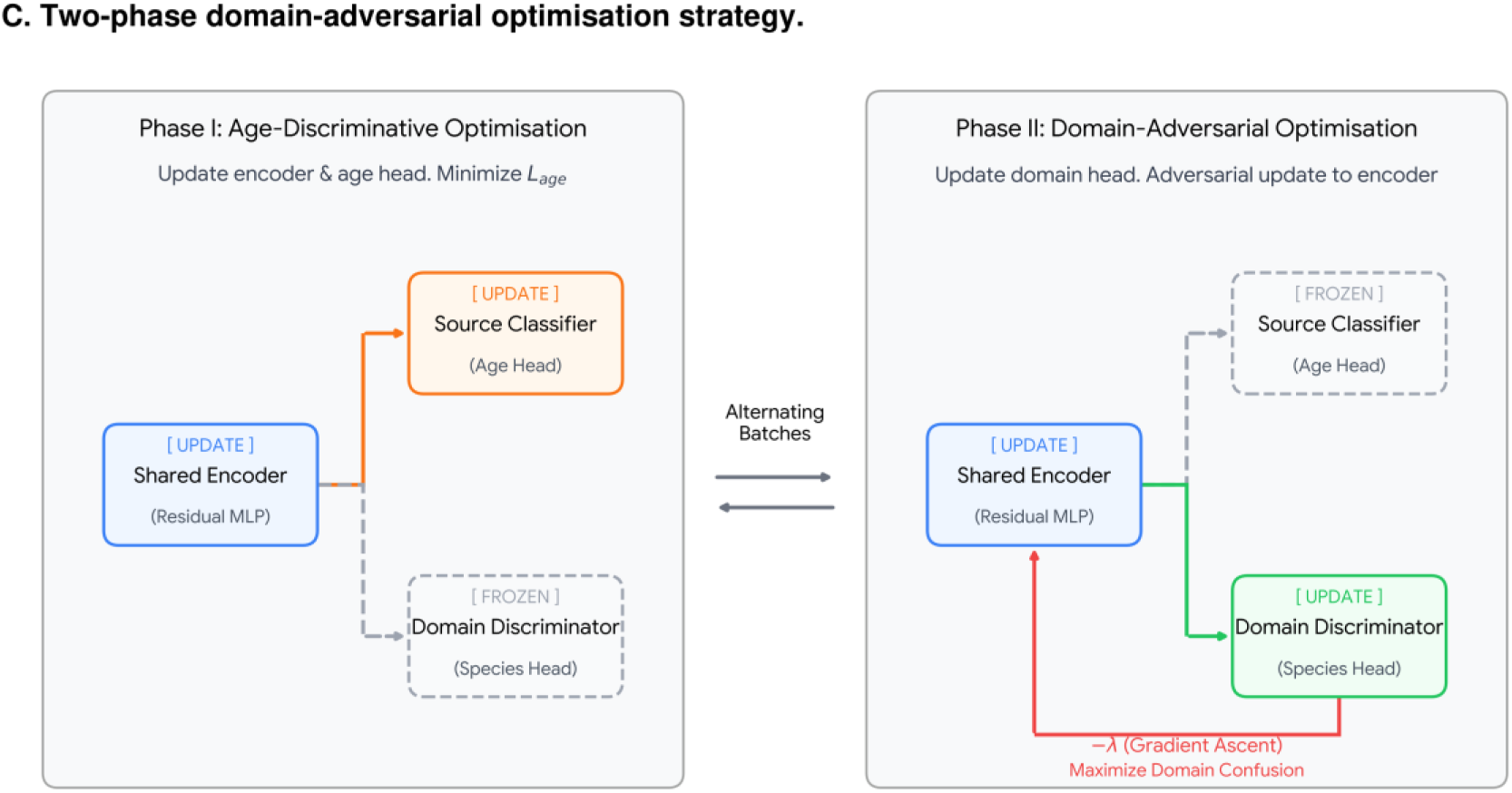
Two-phase residual DANN framework for cross-species HSC ageing-state annotation. a, Overall cross-species annotation pipeline. Human and mouse HSC gene-activity or gene-expression matrices are mapped into a shared latent space and passed to age-classification and species/domain-classification branches. b, Shared feature encoder and multi-branch predictor architecture, including dense layers, batch normalisation, dropout and residual blocks. c, Two-phase adversarial optimisation scheme. In Phase I, the shared encoder and age-classification head are updated while the domain discriminator is frozen. In Phase II, the domain discriminator is updated and the encoder is adversarially adjusted to reduce domain-specific information.

Training diagnostics supported the intended behaviour. Across HSC scRNA-seq, HSC scATAC-seq and CD8+ T-cell scRNA-seq, human-data performance improved across epochs while source and domain losses remained interpretable (Extended Data Figs. 1a,b, 2a,b and 3a,b). Embedding visualisations showed the same transition: early representations were dominated by species separation, whereas later epochs yielded more continuous age-structured manifolds shared between mouse and human cells (Extended Data Figs. 4-6). This progression indicates that the model learns a representation in which cross-species annotation becomes feasible rather than simply fitting mouse labels.

### Cross-species annotation distinguishes HSC ageing states in scRNA-seq and scATAC-seq

We next asked whether mouse-labelled training data could annotate human HSC ageing state in two modalities under a target-validation protocol. Mouse labels supplied the age-classifier gradients, whereas human age labels were used for donor-held validation, checkpoint selection and final reporting. In the HSC scRNA-seq setting, the domain-adversarial model achieved the highest 10-fold cross-validation AUROC among evaluated methods (0.91), exceeding logistic regression (0.58), support vector machine (0.61), random forest (0.61), multilayer perceptron (0.69), ResNet (0.78) and Cellcano (0.71) (Fig. 2a). On the held-out human test set, the final scRNA-seq model reached an AUROC of 0.933, balanced accuracy of 0.897, F1 score of 0.902, sensitivity of 0.889 and specificity of 0.905 (Fig. 2d).

**Figure 2.**
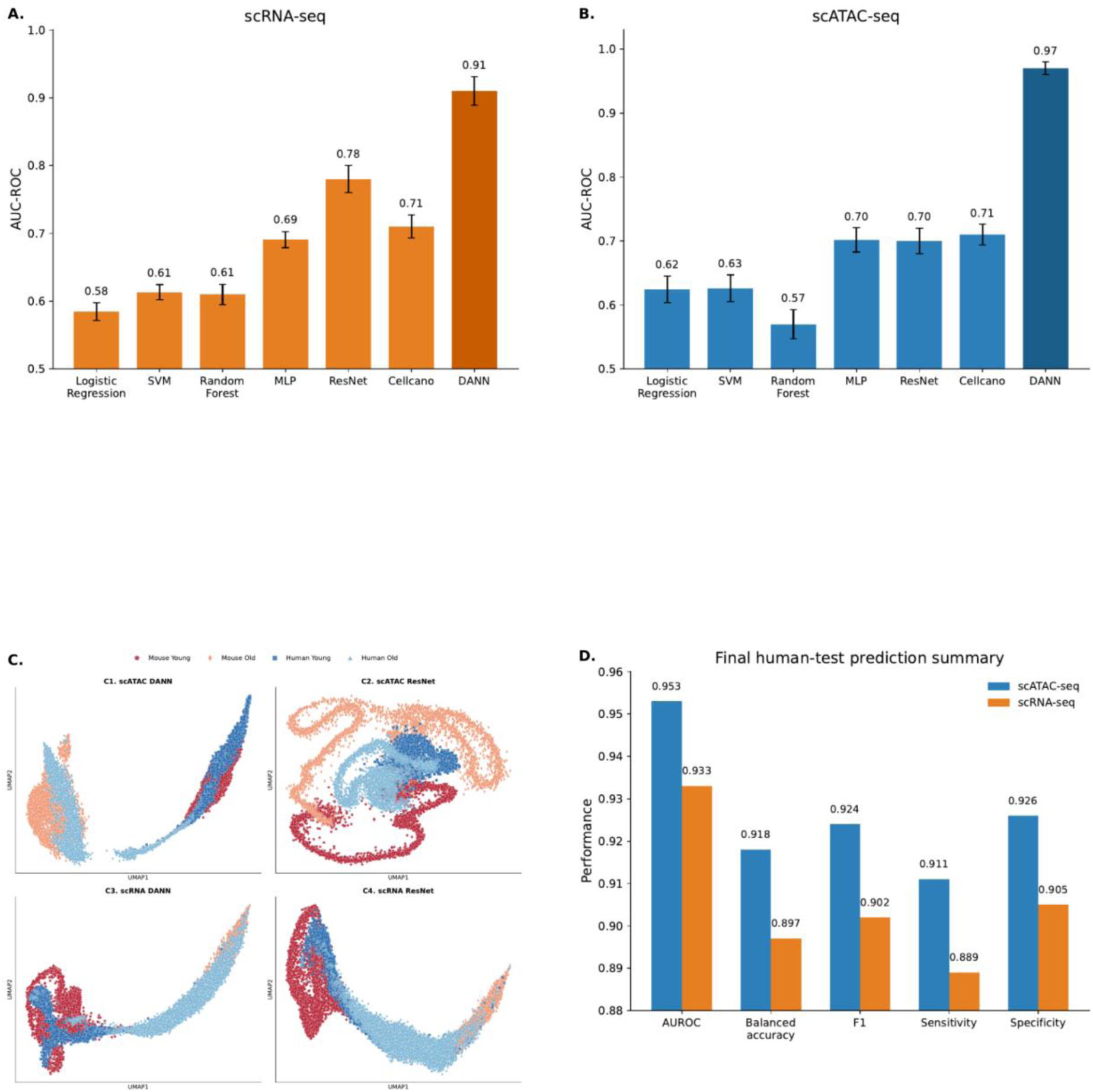
Cross-species annotation of HSC ageing states and comparison with baseline models. a,b, AUROC comparison of baseline classifiers in the HSC scRNA-seq and scATAC-seq settings, including logistic regression, support vector machine (SVM), random forest, multilayer perceptron (MLP), ResNet, Cellcano and DANN. Values were obtained by 10-fold cross-validation; bars show mean AUROC across folds and error bars show s.d. c, UMAP visualisation of latent embeddings learned by DANN and ResNet models in the scATAC-seq and scRNA-seq settings. Cells are coloured by species and age group. d, Final prediction summary on the held-out human test set, showing AUROC, balanced accuracy, F1 score, sensitivity and specificity for the scATAC-seq and scRNA-seq settings.

The advantage over non-adversarial baselines was larger in HSC scATAC-seq. Here the domain-adversarial model reached a cross-validation AUROC of 0.97, whereas the best non-adversarial baselines clustered around 0.70-0.71 (Fig. 2b). Final human-test performance remained high across metrics, with AUROC 0.953, balanced accuracy 0.918, F1 score 0.924, sensitivity 0.911 and specificity 0.926 (Fig. 2d).

UMAP visualisation was consistent with this performance pattern. In both modalities, domain-adversarial embeddings retained age structure while aligning human and mouse cells more effectively than the ResNet baseline, which preserved stronger species segregation (Fig. 2c). These results show that HSC ageing-state annotation is feasible in transcriptomic and chromatin-derived feature spaces when species shift is modelled explicitly.

### Transfer extends to CD8+ T cells and preserves immune-ageing signals

We then tested whether the same framework generalised beyond HSCs and beyond the HSC-centred training context. In an independent cross-species CD8+ T-cell scRNA-seq setting, using the same source-label/target-validation logic, the domain-adversarial model again achieved the best cross-validation AUROC (0.95), exceeding logistic regression (0.62), support vector machine (0.63), random forest (0.57), multilayer perceptron (0.70), ResNet (0.74) and Cellcano (0.72) (Fig. 3a). On the held-out human test set, the final model reached AUROC 0.941, balanced accuracy 0.906, F1 score 0.908, sensitivity 0.926 and specificity 0.885 (Fig. 3b). The learned embedding organised human and mouse CD8+ cells along a shared arch-shaped manifold, whereas the ResNet baseline retained stronger species partitioning (Fig. 3c).

**Figure 3.**
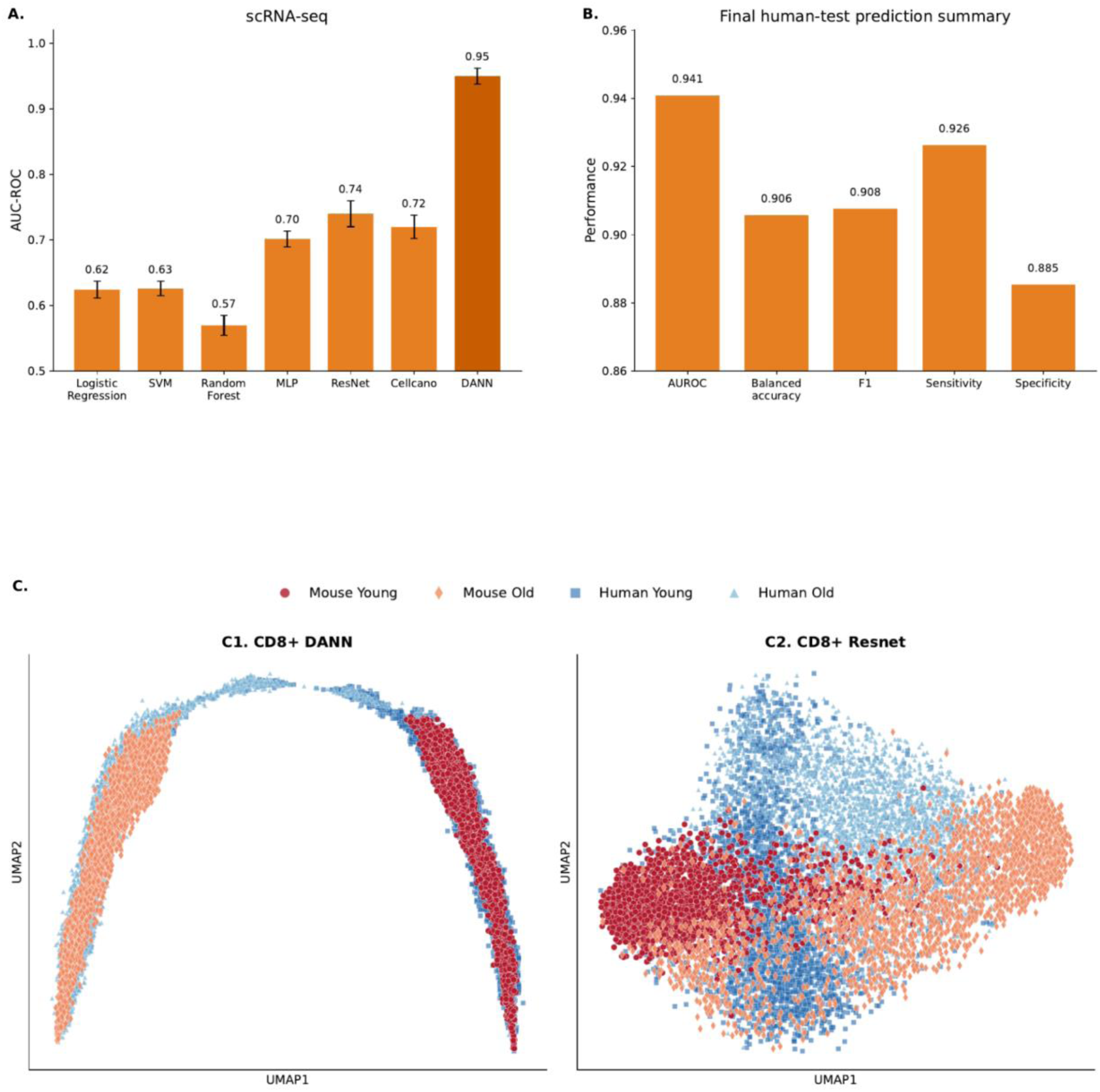
Cross-species CD8+ T-cell ageing-state annotation and comparison with baseline models. a, AUROC comparison of baseline classifiers in the CD8+ T-cell scRNA-seq setting, including logistic regression, support vector machine (SVM), random forest, multilayer perceptron (MLP), ResNet, Cellcano and DANN. Values were obtained by 10-fold cross-validation; bars show mean AUROC across folds and error bars show s.d. b, Final prediction summary on the held-out human test set, showing AUROC, balanced accuracy, F1 score, sensitivity and specificity. c, UMAP visualisation of latent embeddings learned from CD8+ T-cell scRNA-seq data by DANN and ResNet. Cells are coloured by species and age group.

The CD8+ result reduces the likelihood that the HSC performance reflected one dataset or modality alone. The CD8+ setting also supported an independent attribution analysis, in which cross-species signals again included CCL5, CD74, CST7, GZMK and NKG7, and the shared XAI genes were enriched for allograft rejection, TNF-alpha signalling and interferon-response pathways (Extended Data Fig. 9). Thus, the independent CD8+ analysis reproduced both outputs required for the workflow: accurate age-state transfer and conserved immune-ageing features.

### Attribution reveals conserved cross-species ageing features with greater overlap than differential expression

To connect prediction with biological interpretation, we built an attribution workflow around repeated top-K gene selection across DeepLIFT, Integrated Gradients and Saliency (refs. 19–21). Across 190 pairwise run comparisons per method, Integrated Gradients showed the highest top-100 reproducibility (mean Jaccard 0.636 ± 0.037), followed by DeepLIFT (0.585 ± 0.037) and Saliency (0.346 ± 0.031) (Extended Data Fig. 7a). Aggregating repeated runs yielded 824 core genes (frequency ≥90%), 1,165 high-confidence genes (≥80%) and 1,966 medium-confidence genes (≥60%) at the top-3,000 cutoff (Extended Data Fig. 7c).

Against this reproducible background, attribution recovered a different view of conservation from differential expression. The overlap structure in Fig. 4a contained 611 genes captured by cross-species differential expression alone, 265 captured by cross-species XAI alone, and 107 recovered by both approaches. Integrated Gradients further highlighted genes with concordant human and mouse attribution, including SKI, GPR12, FIGN, NOS1AP and KDM6A, while also surfacing genes that were prominent in XAI but not retained by differential-expression ranking (Fig. 4b).

**Figure 4.**
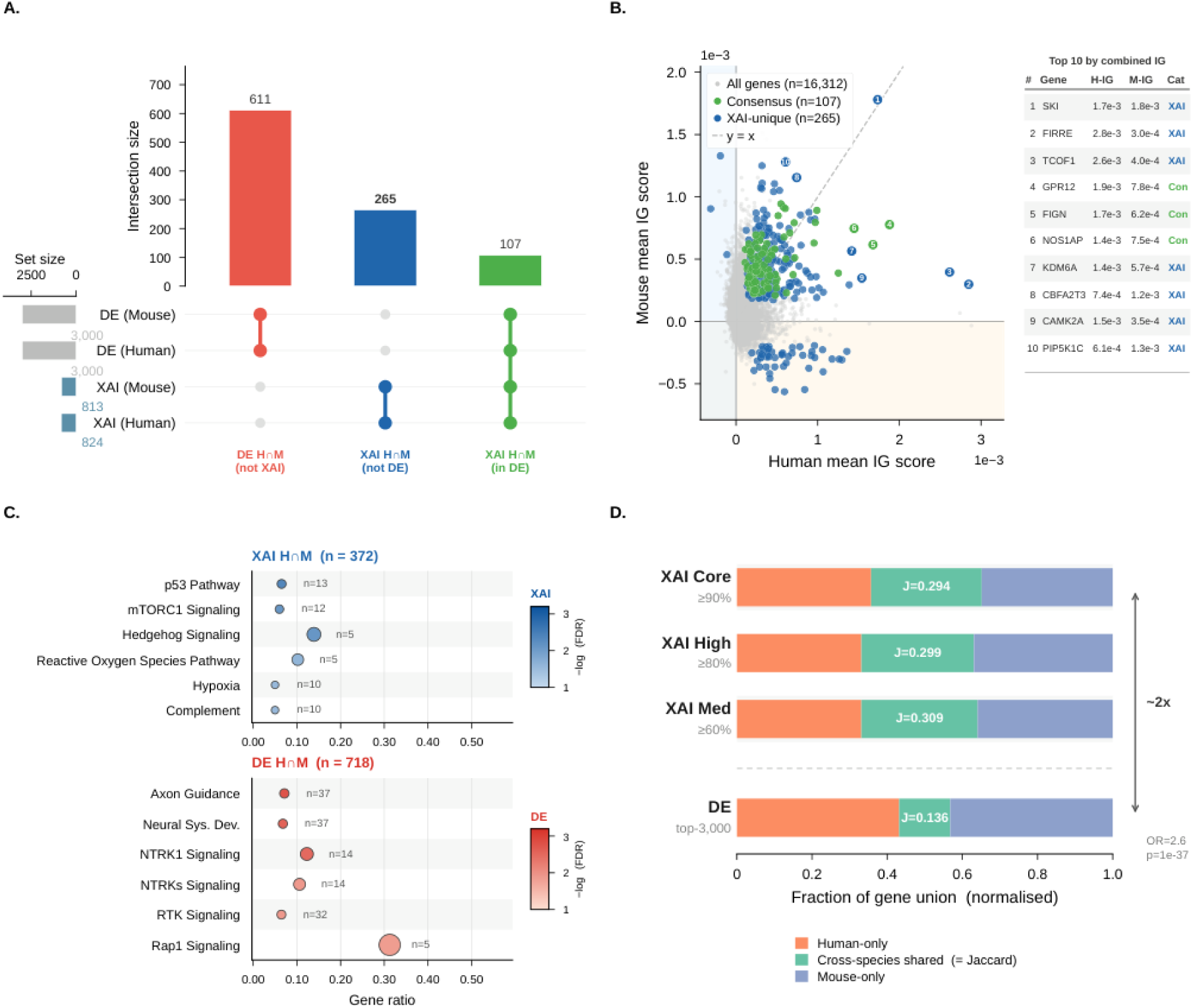
Cross-species comparison of XAI- and differential-expression-derived ageing features. a, UpSet-style summary of overlap between human and mouse gene sets identified by differential expression (DE) and explainable AI (XAI). Numbers indicate the size of each intersection. b, Scatter plot comparing mean integrated gradients (IG) scores in human and mouse, with all genes, consensus genes and XAI-unique genes highlighted. A table lists the top ten genes ranked by combined IG score. c, Pathway enrichment analysis of cross-species shared genes identified by XAI or DE. Dot size indicates gene number and colour indicates enrichment significance. d, Fraction of the gene union assigned to human-only, mouse-only and cross-species shared components for XAI-derived and DE-derived gene sets. Jaccard indices are shown for each gene set.

Pathway-level summaries were compatible with ageing-related processes. Cross-species XAI genes were enriched for p53 pathway, mTORC1 signalling, reactive oxygen species, hypoxia and complement, whereas the differential-expression overlap emphasised pathways such as axon guidance, neural system development, NTRK/RTK signalling and Rap1 signalling (Fig. 4c). The cross-species shared fractions were substantially larger for attribution-derived gene sets than for differential expression alone: the Jaccard indices were 0.294 for the XAI core set, 0.299 for the high-confidence set and 0.309 for the medium-confidence set, compared with 0.136 for the top-3,000 differential-expression set (Fig. 4d). The enrichment of cross-species shared genes was about twofold higher for XAI than for differential expression (odds ratio 2.6, P = 1e-37).

Interpretability also depended on model design. Removing residual connections, replacing ELU with ReLU or reverting to gradient-reversal training reduced predictive performance and weakened attribution stability relative to the full model (Extended Data Fig. 8a-c). These ablations provide empirical support for the residual, ELU-based, two-phase architecture used in the main models.

### Ageing-state annotation of COVID-19 convalescent CD8+ cells associates severe disease with old-like states in younger adults

We next asked whether the learned CD8+ ageing axis could provide a disease-context readout in an external cohort. We applied the pretrained CD8+ model to T3 convalescent samples from the INCOV PBMC scRNA-seq cohort (ArrayExpress E-MTAB-10129; ref. 23), stratified by peak WHO Ordinal Scale (WOS) ≤3 versus 4-7. Each patient was summarised by the fraction of CD8+ cells classified as old-like. Across all ages, the severe group (WOS 4-7, n = 40) followed a distinct age-associated pattern compared with the milder group (WOS ≤3, n = 71), with the largest separation at younger ages (Fig. 5a).

**Figure 5.**
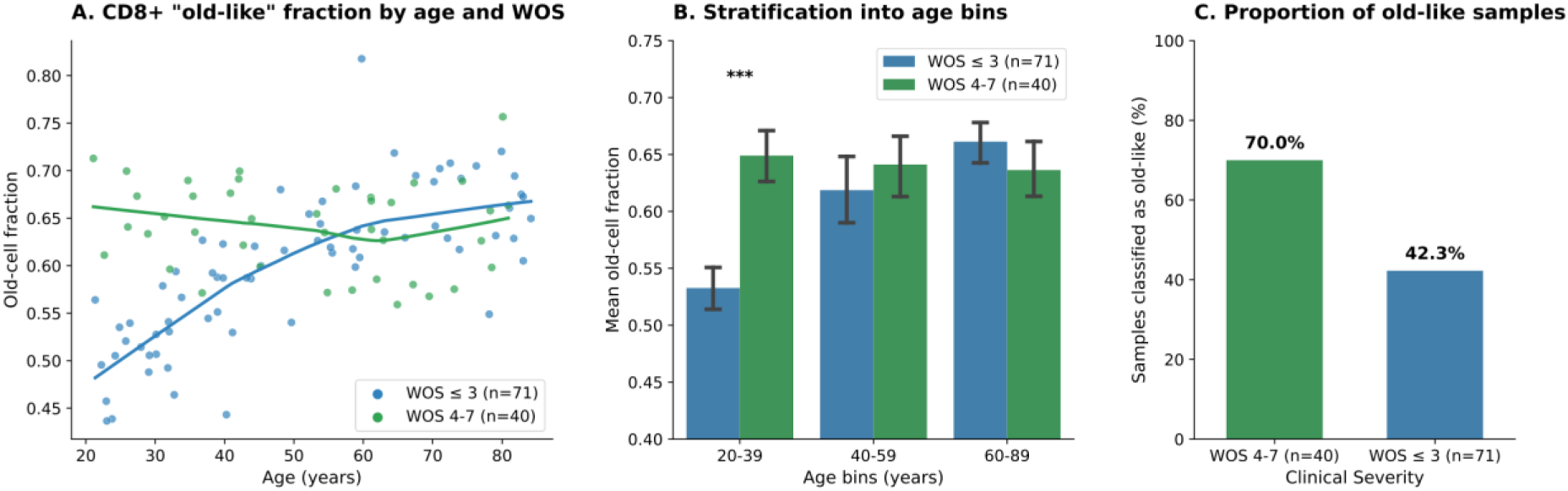
Application of the CD8+ ageing-state model to the COVID-19 cohort. a, Fraction of CD8+ cells classified as old-like plotted against donor age, stratified by peak WHO Ordinal Scale (WOS). Curves show group-wise fitted trends. b, Mean old-cell fraction across age bins (20-39, 40-59 and 60-89 years), stratified by disease severity group. Error bars show 95% bootstrap confidence intervals; bin-wise group differences were assessed with Welch’s t-test and Mann-Whitney U test followed by Benjamini-Hochberg correction, as described in Online Methods. c, Proportion of samples classified as old-like in the two clinical severity groups. Percentages are shown above bars.

Age-binned summaries showed the same separation. In adults aged 20-39 years, the mean old-like CD8+ fraction was markedly higher in the severe group than in the milder group (approximately 0.65 versus 0.53), whereas the difference narrowed in the 40-59 and 60-89 year bins (Fig. 5b). Consistent with this trend, 70.0% of severe cases but only 42.3% of milder cases were classified as old-like at the sample level (Fig. 5c).

This pattern suggests that the learned ageing-state axis captures disease-associated immune variation after infection, particularly in younger adults, in whom severe disease was associated with a larger old-like CD8+ fraction. The analysis is observational and does not establish mechanism, but it illustrates how cross-species annotation can be repurposed as a per-patient immune-state readout.

## Discussion

This study presents a versioned cross-species workflow for ageing-state annotation in haematopoietic stem and immune cells. Mouse-labelled single-cell data supported accurate annotation of held-out human cells across HSC scRNA-seq, HSC scATAC-seq and CD8+ T-cell scRNA-seq when domain shift was modelled explicitly. Combined with attribution and disease application, these analyses connect experimentally tractable animal ageing studies with human single-cell ageing readouts.

The two-phase optimisation scheme improved the behaviour of domain-adversarial learning in sparse single-cell settings. Alternating updates of the age and domain objectives produced stable training curves, consistent embedding trajectories and higher human transfer performance than non-adversarial baselines. The resulting latent spaces reduced species separation while preserving age-related structure, which is the key requirement for cross-species annotation.

Attribution analysis extended the framework from prediction to model-nominated conserved-feature discovery. Repeated attribution across DeepLIFT, Integrated Gradients and Saliency yielded reproducible consensus genes and stronger cross-species overlap than differential expression alone. These findings suggest that multivariate attribution on a transferable model can complement conventional differential analyses when the goal is to prioritise conserved ageing-associated features for follow-up.

A practical advantage of this structure is that prediction, interpretation and release files are linked: the same orthologue-aligned feature space supports model evaluation, attribution and reviewer reruns. The resource therefore provides a reproducible path from animal-labelled experiments to human single-cell ageing readouts, rather than a classifier benchmark alone.

The COVID-19 analysis extends the workflow to disease by summarising each patient as the fraction of CD8+ cells classified as old-like. This readout was associated with disease severity primarily in younger adults, suggesting that the learned ageing axis captures disease-associated immune variation after infection. Several limitations remain. Cross-species alignment is designed to suppress domain-specific signal and may therefore underrepresent genuinely species-restricted ageing programmes. Similarly, scATAC-derived gene activity is a gene-level approximation, and distal regulatory effects or niche-derived signals are represented only when they leave measurable cell-intrinsic profiles. The COVID-19 analysis is observational, and attribution nominates candidate regulators rather than establishing causality. Functional perturbation studies and additional human datasets will be required to define the mechanistic basis and generalisability of these associations.

## Online Methods

### Study design and data sources

The study was designed to test whether ageing-state labels learned from mouse single-cell data could be transferred to human cells while preserving interpretability. The main analyses were performed in HSCs, in which mouse cells provided source-domain age labels and human cells formed the target domain. Human age labels were withheld from gradient-based model fitting and used only for validation, checkpoint selection and final reporting. The same workflow was then evaluated in an independent CD8+ T-cell scRNA-seq setting and applied to a COVID-19 convalescent cohort.

For HSC scATAC-seq, mouse bone marrow HSCs from 10 eight-week mice and 2 twenty-four-month mice were profiled with the optimised DOGMA-seq/10x Multiome workflow (ref. 24) and processed with Cell Ranger ARC v2.0.0 and ArchR v1.0.1 (ref. 25). The cross-species HSC resource also included young and old human HSC scATAC-seq datasets processed through the same Cell Ranger ARC/ArchR pipeline. Barcode-level quality control retained human cells with at least 3,000 unique fragments and transcription start site (TSS) enrichment of at least 10, and mouse cells with at least 2,500 unique fragments and TSS enrichment of at least 8. Gene-score matrices were generated in ArchR, restricted to one-to-one human-mouse orthologues from Ensembl BioMart release 98 (ref. 26), and log-transformed, quantile-normalised and standardised before modelling. This yielded 16,312 shared features across 8,026 human HSCs (2,974 young and 5,052 old) and 7,218 mouse HSCs (3,013 young and 4,205 old). Human target data were partitioned at the donor level so that the final held-out human test set remained fully independent of model development.

For HSC scRNA-seq, gene-expression matrices were derived from the matched gene-expression modality of the same cross-species HSC resource and processed under the same Cell Ranger-based workflow, quality-control logic and orthologue-mapping strategy used for the scATAC-seq analysis. After quality control, the scRNA-seq dataset comprised 3,012 young and 3,723 old mouse HSCs, and 2,177 young and 5,034 old human HSCs. Orthologue-aligned expression matrices were log-normalised and standardised before model fitting, and the scRNA-seq analysis used the same model architecture, baseline set and donor-aware evaluation framework as the HSC scATAC-seq analysis.

External validation in CD8+ T cells used public mouse and human scRNA-seq datasets. Mouse CD8+ cells were obtained from spleen, peritoneum, lung and liver of young (3-4 months) and aged (24 months) male C57BL/6 mice through Single Cell Explorer (ref. 5). Human conventional CD8+ T cells were obtained from the Single-Cell Atlas of Human Blood During Healthy Aging (Synapse syn49637038; ref. 6), comprising samples from healthy donors aged 25-85 years. Mouse cells with fewer than 500 detected genes or more than 10% mitochondrial UMIs were removed, and genes detected in fewer than 5 cells were excluded. Human cells with fewer than 600 detected genes or more than 15% mitochondrial transcripts were removed, and genes expressed in fewer than 10 cells were excluded. In both species, counts were normalised as log(counts per 10,000 + 1), the top 2,000 highly variable genes were identified in Seurat v3.2 (ref. 8), and PCA, graph-based clustering and UMAP were used for quality control. For cross-species transfer, up to 3,000 young and 3,000 old cells per species were subsampled and restricted to 16,041 one-to-one orthologues.

### Feature construction and orthologue mapping

For scATAC-seq, gene scores were generated in ArchR after barcode-level filtering. For scRNA-seq, orthologue-aligned gene-expression matrices were log-normalised before standardisation. In all settings, mouse-labelled cells formed the source domain and human cells formed the target domain; target-domain age labels were not used in the adversarial loss or age-classifier gradient updates. Features were standardised to zero mean and unit variance before model fitting, and feature order, scaling choices and class order were recorded with the corresponding released models and reproduction manifests. For the COVID-19 application, genes absent after alignment to the reference CD8+ feature list were set to zero.

### Model architecture and two-phase domain-adversarial optimisation

The model consisted of a shared residual multilayer perceptron encoder followed by two heads: an age classifier and a species discriminator (Fig. 1b). The encoder comprised a 2,048-node input projection, three residual fully connected blocks and a 512-node bottleneck, with batch normalisation, ELU activation and dropout throughout. The age head and domain head each contained a 512-node hidden layer followed by a two-class softmax output. The classifier head predicted young versus old, and the discriminator predicted mouse versus human.

Let *x_i_* denote the orthologue-aligned feature vector for cell *i*. Source cells have age labels *y_i_* ∈ {young, old}, and each cell has a species/domain label *z_i_* ∈ {mouse, human}. The shared encoder, age head and domain head are denoted by *f*, *g* and *h*, respectively. For mouse-labelled source cells, the age loss was

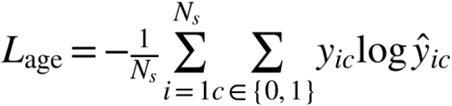

For mixed human-mouse batches, the domain loss was

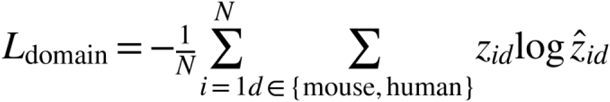

A joint adversarial objective can be written as

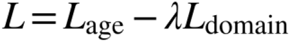

where λ controls the strength of domain confusion.

In Phase I, the encoder and age head were updated while the domain discriminator was frozen,

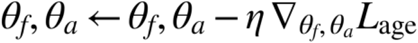

In Phase II, the domain head was updated to minimise domain loss and the encoder was adversarially adjusted to maximise it,

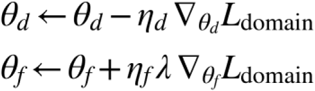

This alternating optimisation approximates block-coordinate descent/ascent and was used instead of a gradient-reversal layer (ref. 14). All DANN models used AdamW (refs. 27,28) with exponential learning-rate decay, clipnorm 1.0, *β_2_* = 0.999 and *ε* = 1e-7; feature-extractor updates used *β_1_* = 0.96, whereas classifier-head updates used *β_1_* = 0.9. For HSC scATAC-seq, initial learning rates were 1.0e-3 for the encoder and domain head and 1.2e-4 for the age head. For HSC scRNA-seq, the corresponding initial learning rates were 1.2e-4, 1.2e-4 and 1.3e-5, respectively. For CD8+ T-cell scRNA-seq, the encoder and domain head used 1.5e-3 and the age head used 1.5e-5. Mini-batches contained 512 cells, balanced between mouse and human domains. Checkpoint selection used target-domain validation accuracy where human age labels were available; final human-test summaries were evaluated after checkpoint selection with patience windows of 200, 100 and 300 iterations for HSC scATAC-seq, HSC scRNA-seq and CD8+ T-cell models, respectively.

### Baseline models and evaluation

The domain-adversarial model was compared against logistic regression, support vector machine, random forest, multilayer perceptron, ResNet and Cellcano baselines. Cellcano was implemented according to its published two-round supervised annotation workflow (ref. 29). In brief, a first-round multilayer perceptron was trained on the source data, prediction entropy was used to select confident target anchors, and a second-round knowledge-distillation model was trained on these anchors to refine target predictions. We applied this workflow to the same orthologue-aligned gene-level matrices used in the transfer setting. Classical models were tuned by grid search on mouse training data and evaluated with the same cross-species splits. Except for the final held-out human test summaries, all reported model-comparison metrics were aggregated across 10-fold cross-validation. Performance was summarised using AUROC and, for final held-out human tests, balanced accuracy, F1 score, sensitivity and specificity. UMAP was used to visualise latent structure from DANN and ResNet embeddings (ref. 30).

### Attribution analysis and consensus gene selection

Attribution was used to identify genes that pushed model outputs towards the old-state label. We computed DeepLIFT, Integrated Gradients and Saliency scores using Captum (ref. 22). For each method, attribution was repeated across 20 runs, the top 3,000 genes per run were retained, and consensus frequency was calculated across runs and species. Genes present in at least 90%, 80% and 60% of runs were labelled as core, high-confidence and medium-confidence, respectively. Reproducibility was quantified by pairwise Jaccard similarity of the top 100 genes across the 190 run pairs per method, and concordance across methods was assessed over a range of top-N cutoffs, as shown in Extended Data Fig. 7.

### Differential expression and pathway enrichment

Differential expression was used as a comparator rather than the primary discovery engine. Human and mouse age contrasts were analysed separately with MAST, cross-species overlaps were constructed by intersecting the resulting gene sets across species, and pathway enrichment was performed with MsigDB Hallmark gene sets (refs. 31,32). The shared and species-specific fractions of XAI- and DE-derived gene sets were quantified by Jaccard similarity and enrichment statistics.

### COVID-19 application analysis and statistics

For the COVID-19 cohort, the pretrained CD8+ model was applied at single-cell resolution after feature alignment. A cell was counted as old-like only when it was assigned to the old class with maximum class probability of at least 0.80; each patient was then summarised by the old-like fraction

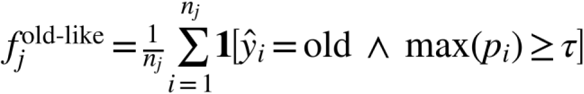

Here *n_j_* denotes the number of retained CD8+ cells in patient *j*, and the confidence threshold was fixed at *τ* = 0.80. LOWESS smoothing (ref. 33) was used for visualisation with an adaptive span frac = clip(0.75 - 0.003(n - 10), 0.45, 0.75) and it = 1, with 95% bootstrap confidence intervals from 500 resamples. For age-binned analyses, ages were partitioned into 20-39, 40-59 and 60-89 years, and per-bin means were reported with 95% bootstrap confidence intervals from 2,000 resamples (seed 2024). Differences between WOS ≤3 and WOS 4-7 groups were tested within each bin by Welch’s t-test and Mann-Whitney U test, followed by Benjamini-Hochberg correction across bins.

## Ethics and consent

All analyses used de-identified single-cell datasets and processed matrices; no new human participants were recruited for this computational study. For the HSC datasets, human bone marrow data were generated under the original study approvals. Older-donor bone marrow was obtained after informed consent through the Mechanisms of Age-Related Clonal Haematopoiesis (MARCH) Study (REC Ref: 17/YH/0382), and young-donor bone marrow was obtained from commercial suppliers. The mouse experiments associated with the HSC resource were performed according to UK Home Office regulations and with ethical approval from the University of Oxford Medical Sciences Division Animal Welfare and Ethical Review Board. Public CD8+

T-cell and COVID-19 datasets were analysed in de-identified or aggregate form under their original study approvals and data-access terms.

## Data availability

Processed matrices, metadata tables, feature lists and trained model weights have been deposited on Hugging Face Hub at https://huggingface.co/datasets/simiaoAA/cross-species-aging-data and https://huggingface.co/simiaoAA/cross-species-aging-models. Public source datasets used for external validation and clinical application are available through Synapse (syn49637038), ArrayExpress (E-MTAB-10129) and the source accessions described in Online Methods.

## Code availability

Source code, environment files, input manifests and reviewer-facing reproduction instructions are available at https://github.com/SimiaoZhao/cross-species-aging-resource. The release includes scripts to download artifacts, verify required inputs, sanity-check HSC scRNA young/old labels, evaluate released DANN models and run from-scratch HSC scRNA DANN training with recorded seeds and hyperparameters.

## Supporting information

Extended Data Fig. 4. HSC scATAC-seq DANN embedding evolution

Extended Data Fig. 5. HSC scRNA-seq DANN embedding evolution

Extended Data Fig. 6. CD8+ T-cell scRNA-seq DANN embedding evolution

Extended Data Fig. 7. Attribution reproducibility and consensus analysis

Extended Data Fig. 8. Model ablation and architecture sensitivity

Extended Data Fig. 9. CD8+ T-cell cross-species attribution analysis

Extended Data Fig. 1. HSC scRNA-seq training curves

Extended Data Fig. 2. HSC scATAC-seq training curves

Extended Data Fig. 3. CD8+ T-cell scRNA-seq training curves

---

**Extended Data Fig. 1.**
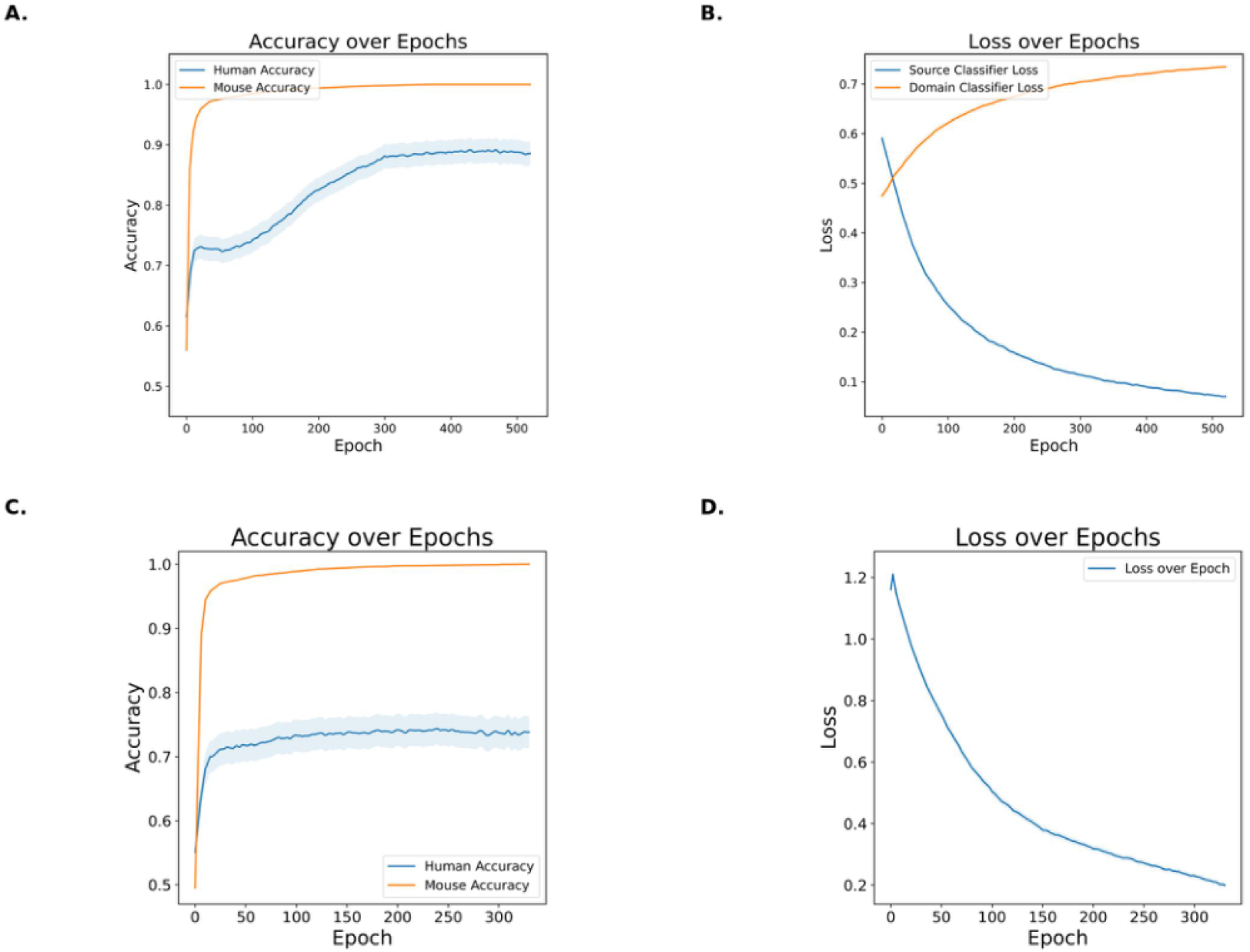
Training curves for HSC scRNA-seq models under 10-fold cross-validation. a, Prediction accuracy over training epochs for the DANN model, shown separately for human and mouse data. b, Source-classifier loss and domain-classifier loss over training epochs for the DANN model. c, Prediction accuracy over training epochs for the ResNet baseline, shown separately for human and mouse data. d, Training loss over epochs for the ResNet baseline. For each epoch, values summarise results aggregated across the ten cross-validation folds.

**Extended Data Fig. 2.**
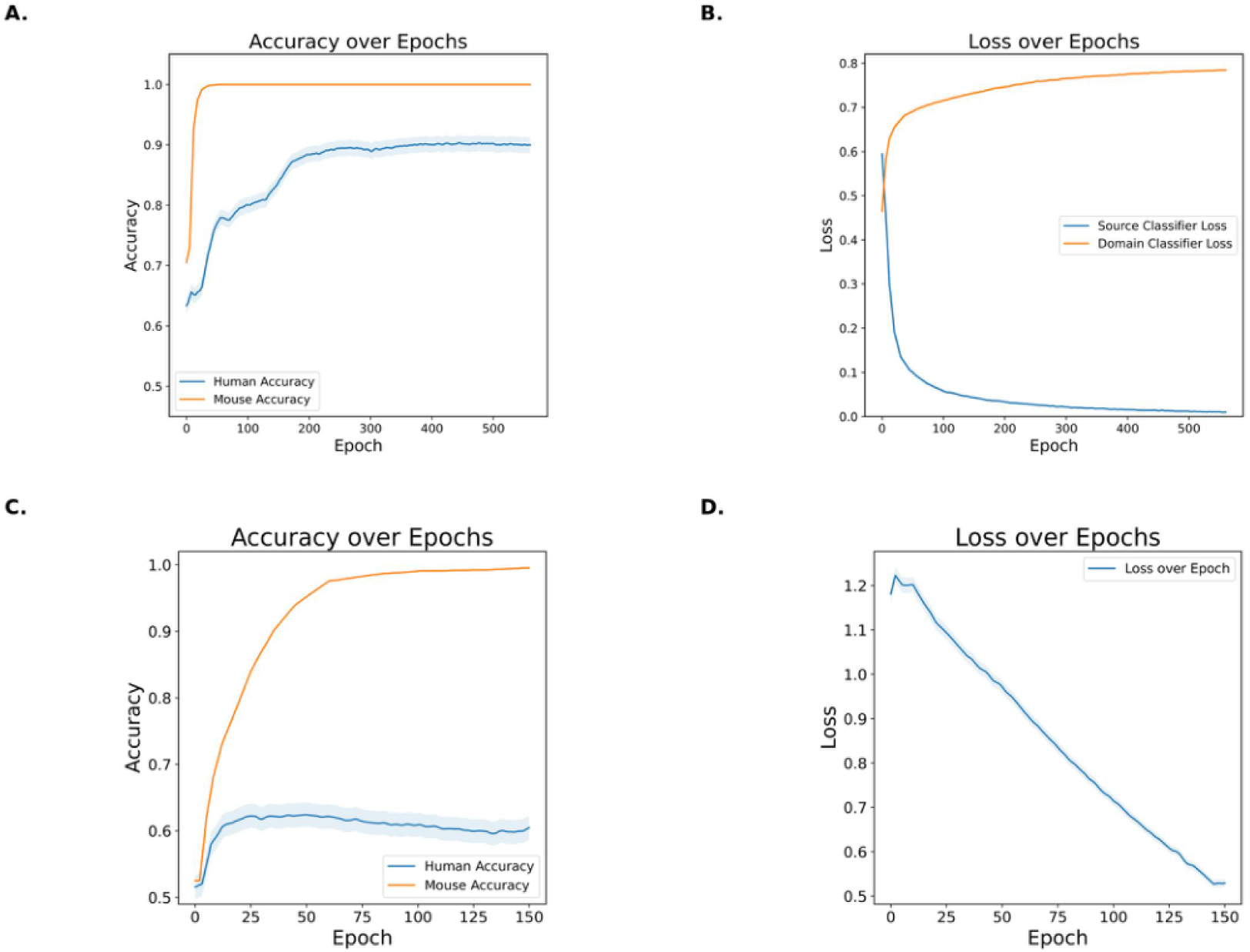
Training curves for HSC scATAC-seq models under 10-fold cross-validation. a, Prediction accuracy over training epochs for the DANN model, shown separately for human and mouse data. b, Source-classifier loss and domain-classifier loss over training epochs for the DANN model. c, Prediction accuracy over training epochs for the ResNet baseline, shown separately for human and mouse data. d, Training loss over epochs for the ResNet baseline. For each epoch, values summarise results aggregated across the ten cross-validation folds.

**Extended Data Fig. 3.**
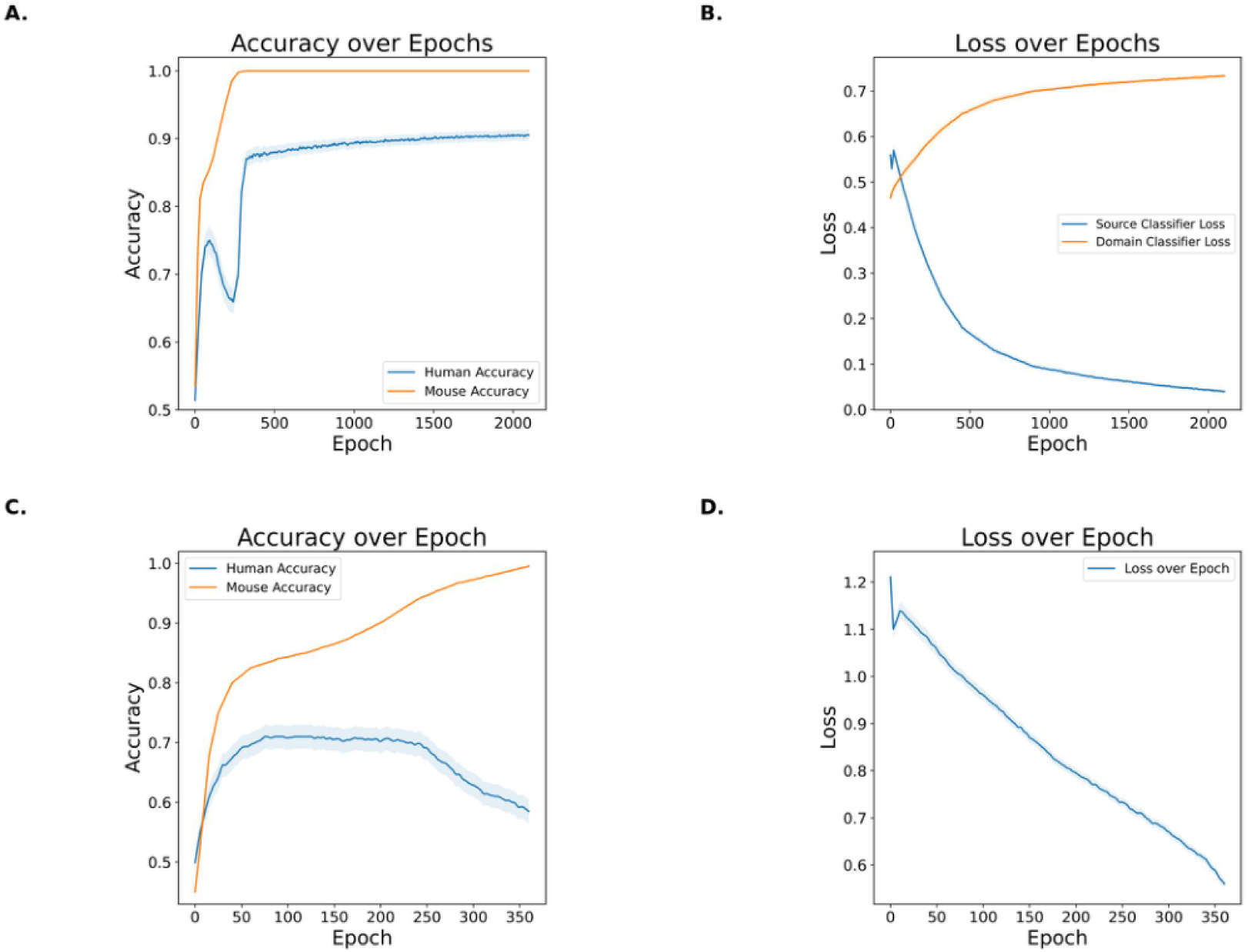
Training curves for CD8+ T-cell scRNA-seq models. a, Prediction accuracy over training epochs for the DANN model, shown separately for human and mouse data. b, Source-classifier loss and domain-classifier loss over training epochs for the DANN model. c, Prediction accuracy over training epochs for the ResNet baseline, shown separately for human and mouse data. d, Training loss over epochs for the ResNet baseline. For each epoch, values summarise results aggregated across the ten cross-validation folds.

**Extended Data Fig. 4.**
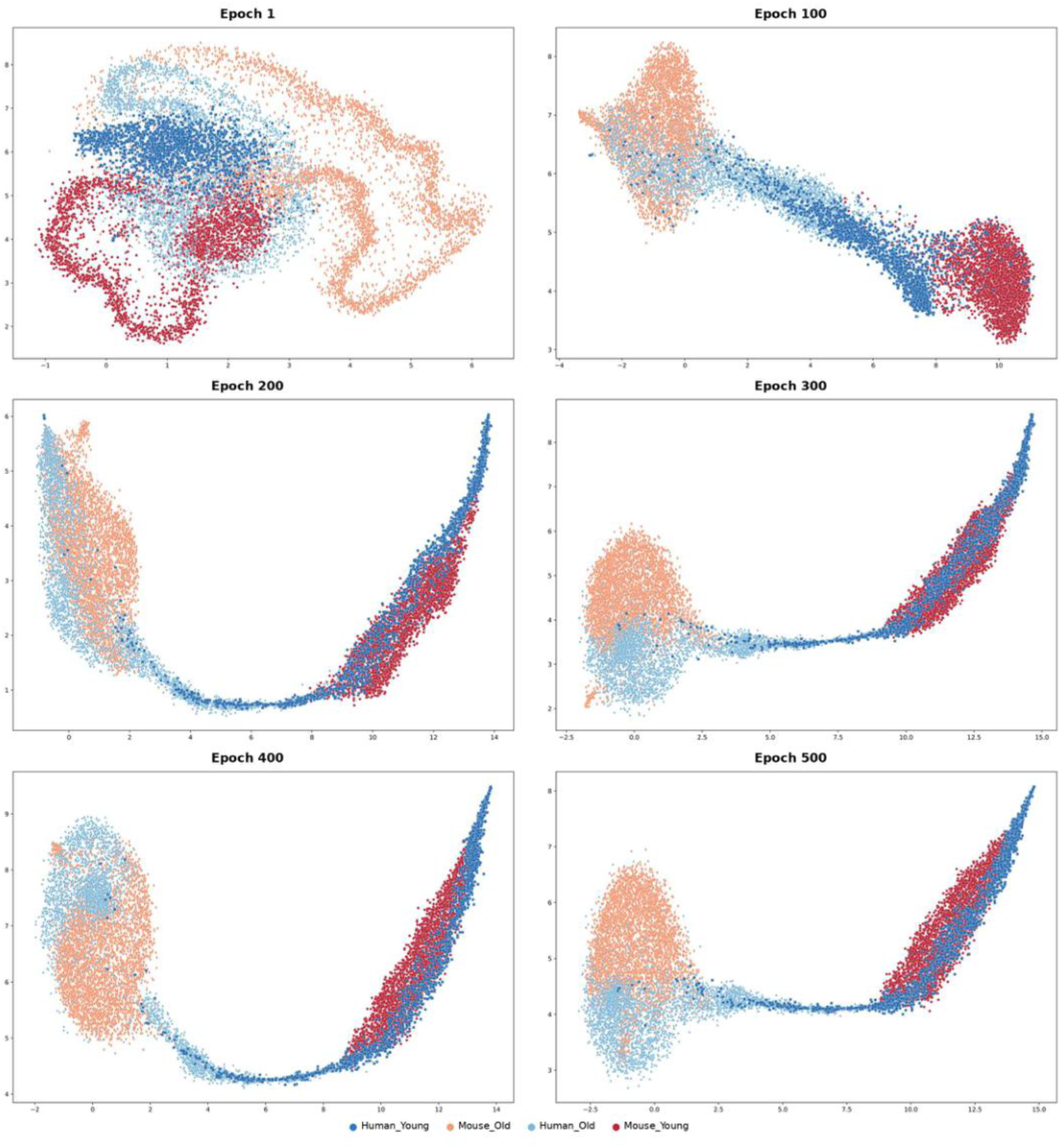
Evolution of HSC scATAC-seq DANN embeddings during training. UMAP visualisation of latent embeddings from the HSC scATAC-seq DANN model at epochs 1, 100, 200, 300, 400 and 500. Cells are coloured by species and age group: human young, human old, mouse young and mouse old.

**Extended Data Fig. 5.**
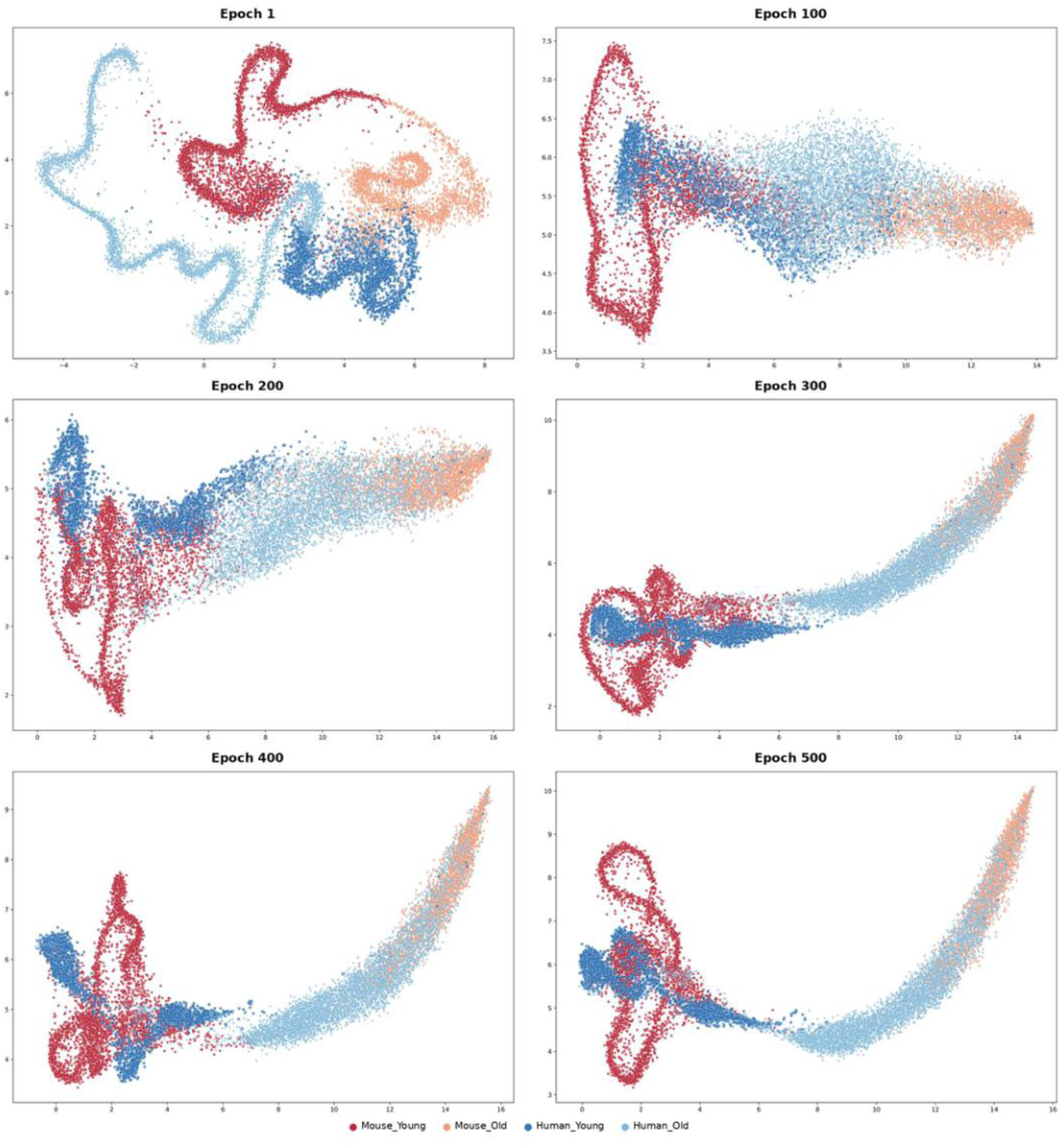
Evolution of HSC scRNA-seq DANN embeddings during training. UMAP visualisation of latent embeddings from the HSC scRNA-seq DANN model at epochs 1, 100, 200, 300, 400 and 500. Cells are coloured by species and age group: human young, human old, mouse young and mouse old.

**Extended Data Fig. 6.**
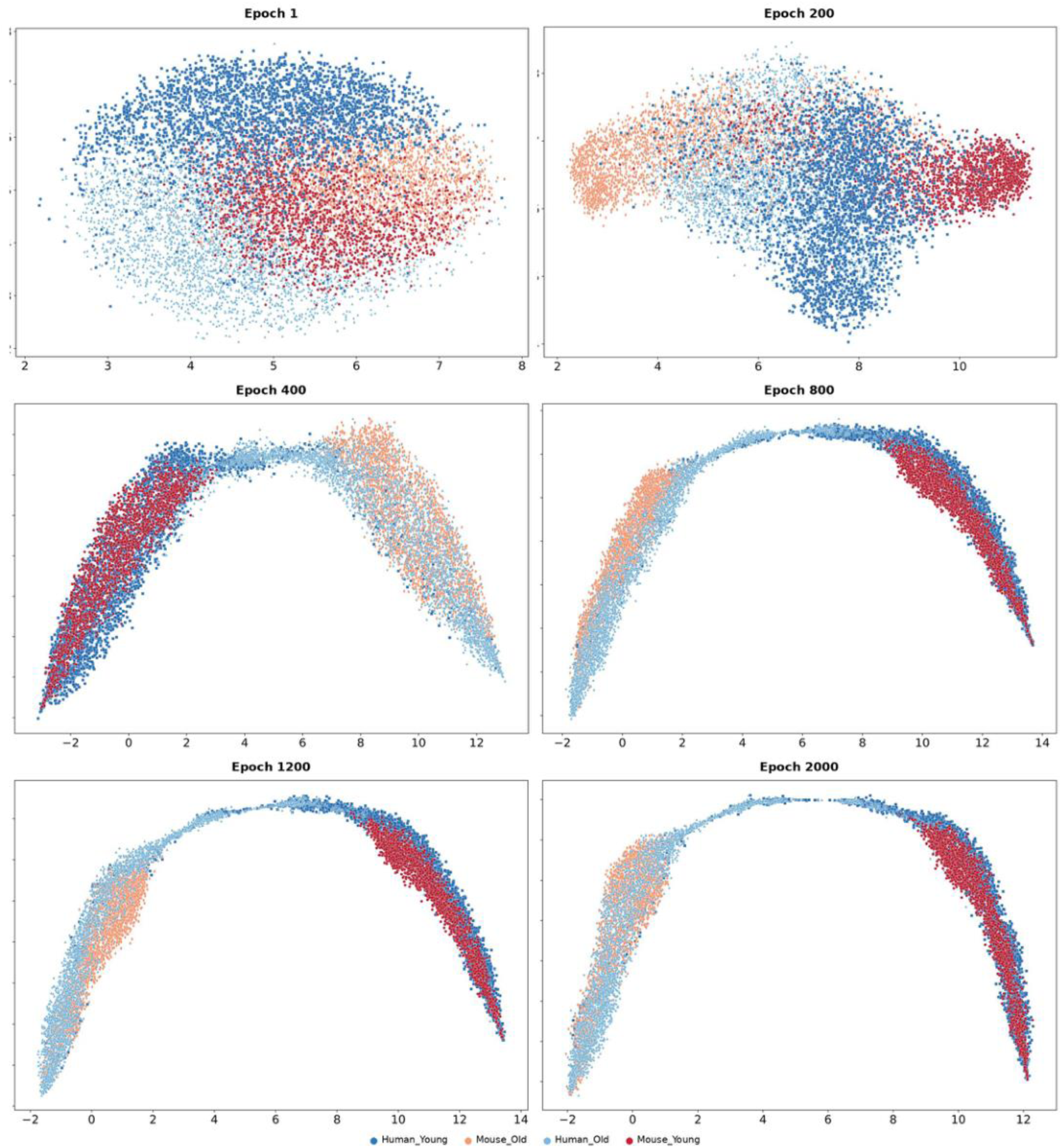
Evolution of CD8+ T-cell scRNA-seq DANN embeddings during training. UMAP visualisation of latent embeddings from the CD8+ T-cell scRNA-seq DANN model at epochs 1, 200, 400, 800, 1200 and 2000. Cells are coloured by species and age group: human young, human old, mouse young and mouse old.

**Extended Data Fig. 7.**
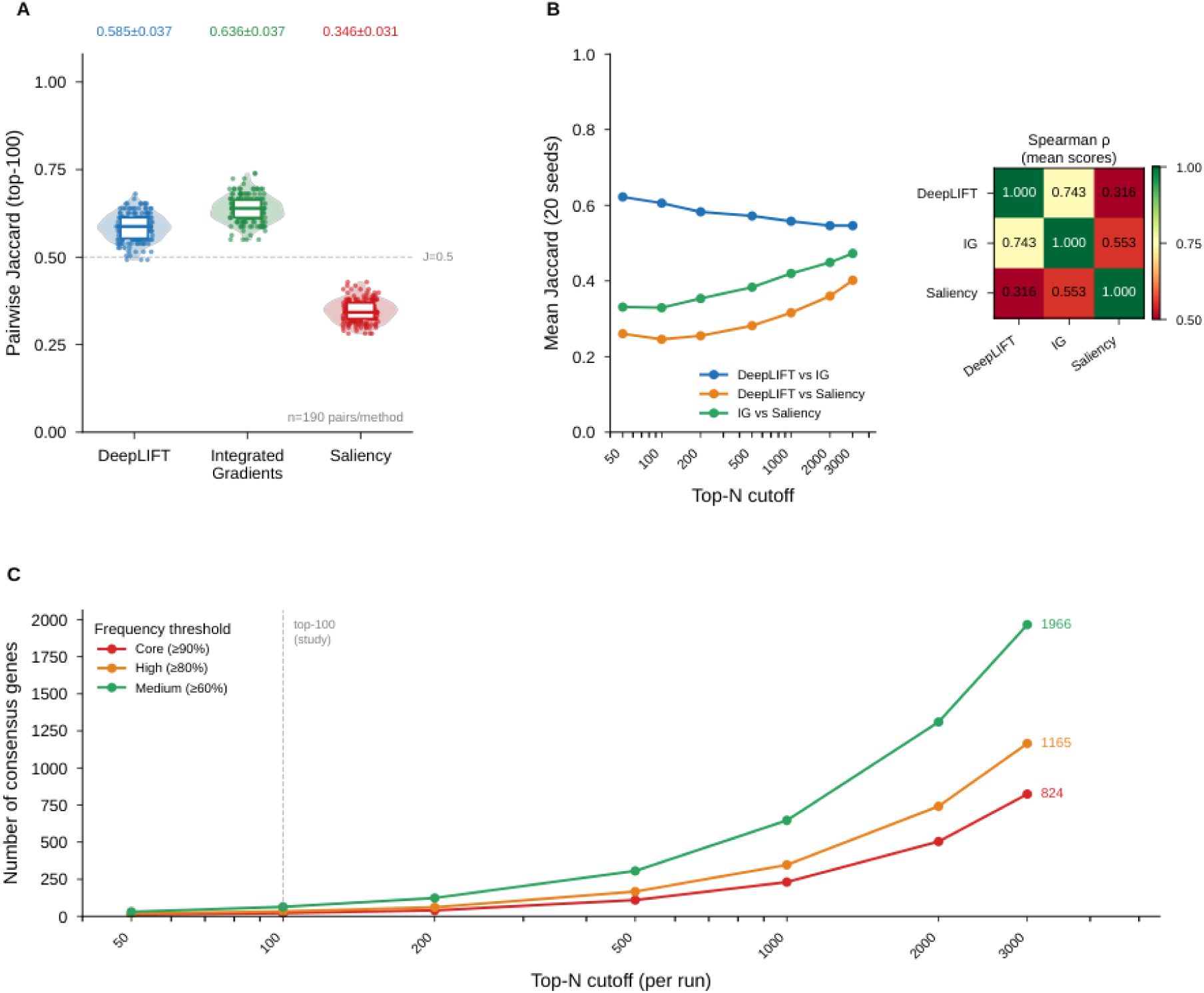
Reproducibility and consensus analysis of attribution methods. a, Pairwise Jaccard similarity of top-100 genes identified across repeated runs using DeepLIFT, Integrated Gradients and Saliency. Values are shown as mean ± s.d. b, Mean Jaccard similarity across different top-N cutoffs for pairwise comparisons between attribution methods. A heatmap shows Spearman correlation coefficients between mean attribution scores from the three methods. c, Number of consensus genes identified across repeated runs at different top-N cutoffs under three frequency thresholds: core (≥90%), high (≥80%) and medium (≥60%).

**Extended Data Fig. 8.**
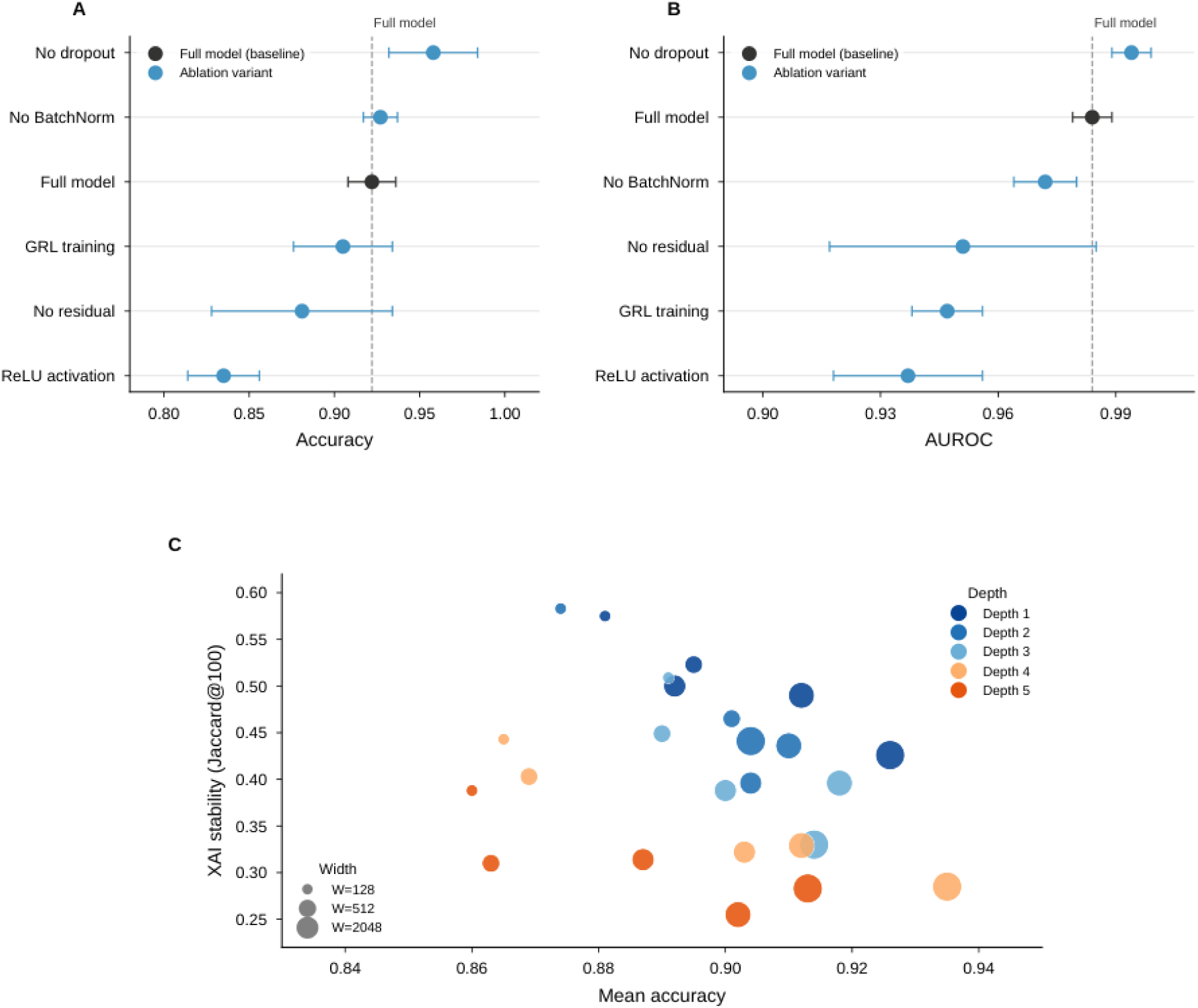
Ablation and architecture-sensitivity analysis of the model. a, Accuracy of the full model and ablation variants, including no dropout, no batch normalisation, no residual connections, GRL training and ReLU activation. b, AUROC of the full model and ablation variants shown in a. c, Relationship between mean accuracy and XAI stability (Jaccard@100) across models with different depths and widths. Point colour indicates network depth and point size indicates network width. All variants were evaluated with the same data splits and reporting metrics used for the corresponding main models.

**Extended Data Fig. 9.**
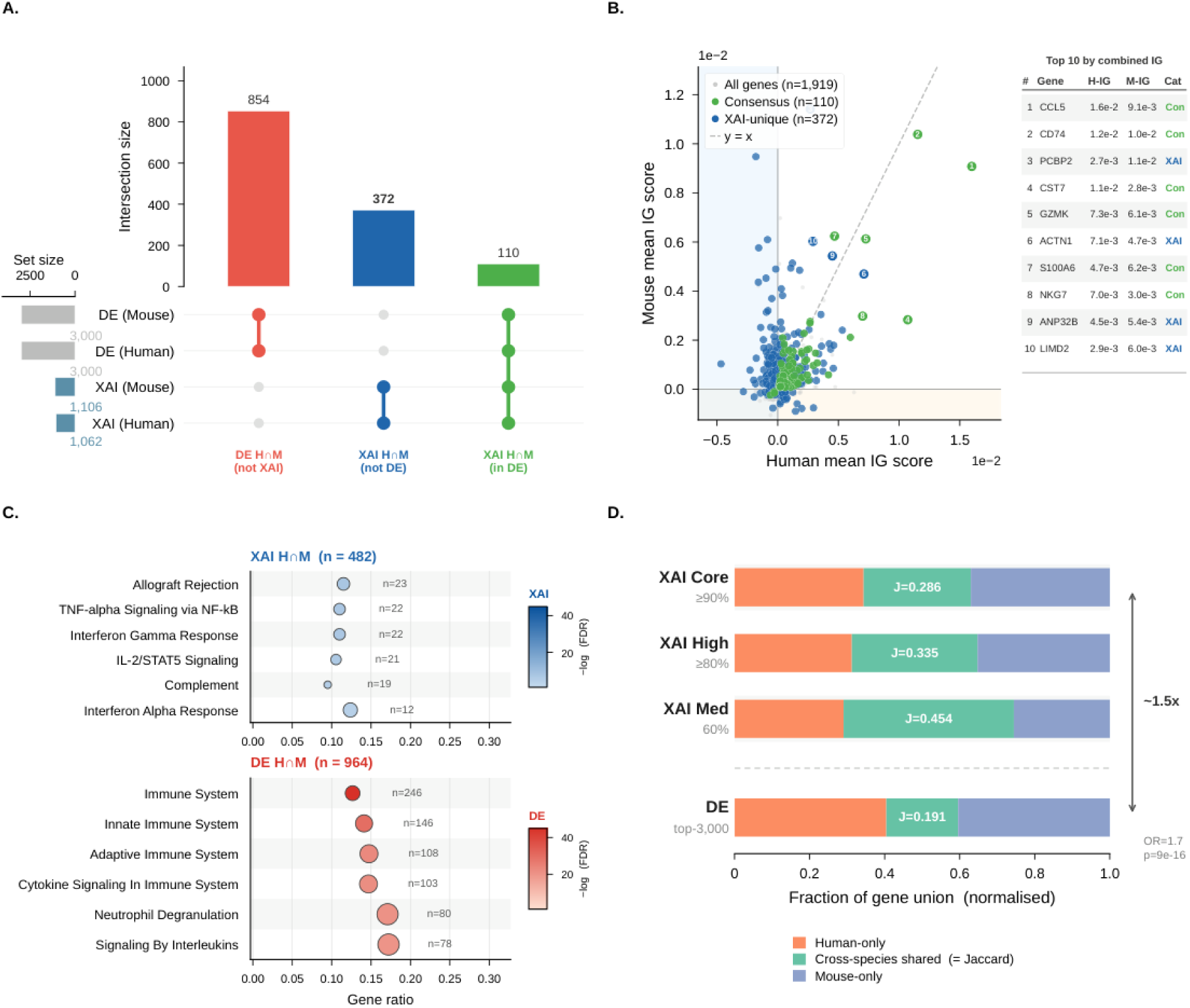
Cross-species attribution analysis in CD8+ T cells. a, UpSet-style summary of overlap between human and mouse gene sets identified by differential expression (DE) and explainable AI (XAI). Numbers indicate the size of each intersection. b, Scatter plot comparing mean integrated gradients (IG) scores in human and mouse, with all genes, consensus genes and XAI-unique genes highlighted. A table lists the top ten genes ranked by combined IG score. c, Pathway enrichment analysis of cross-species shared genes identified by XAI or DE. Dot size indicates gene number and colour indicates enrichment significance. d, Fraction of the gene union assigned to human-only, mouse-only and cross-species shared components for XAI-derived and DE-derived gene sets. Jaccard indices are shown for each gene set.

