## Supplementary figures and images for "Deep learning enables cross-species annotation and attribution of ageing states in haematopoietic stem and immune cells"

### Extended Data Fig. 1. HSC scRNA-seq training curves

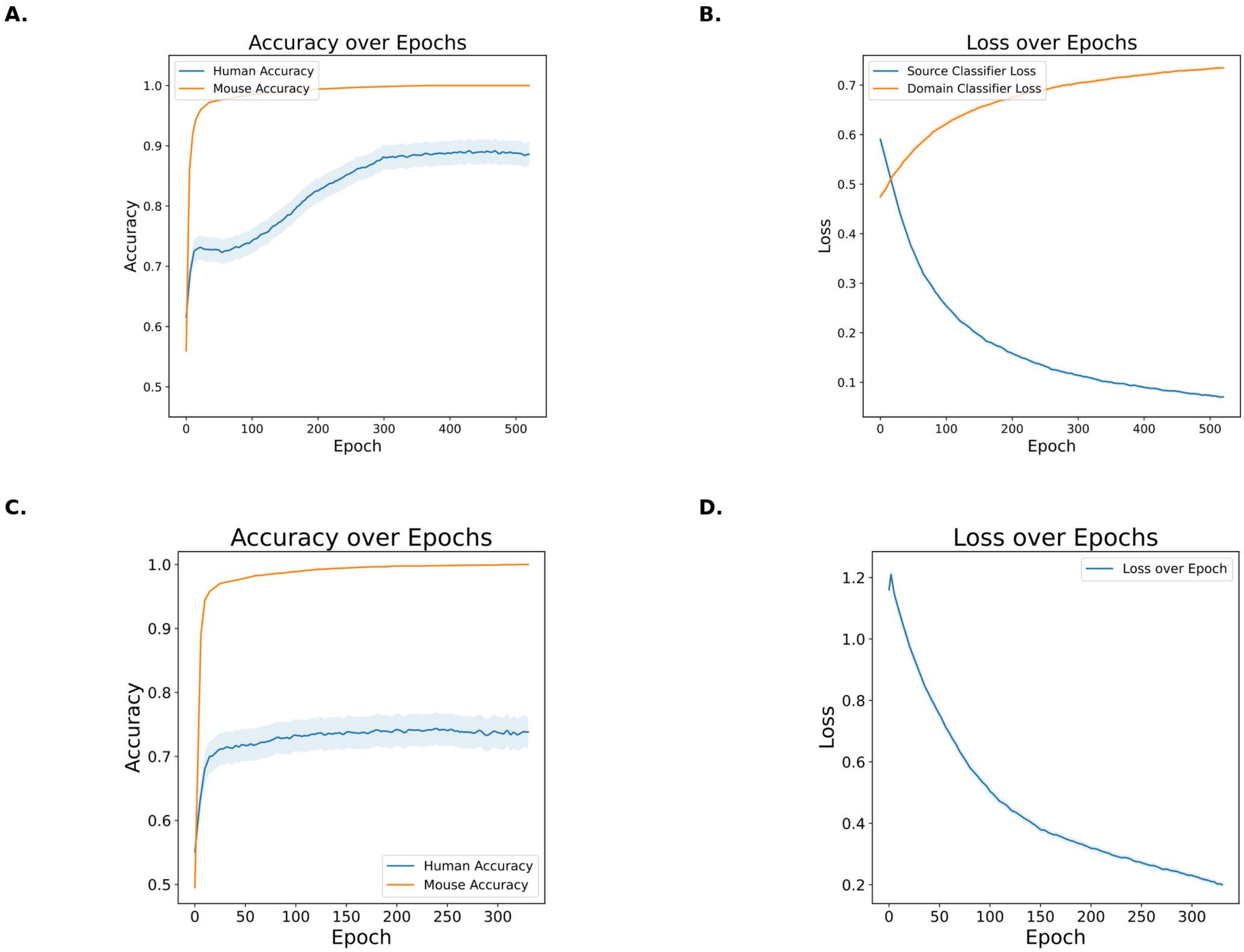

### Extended Data Fig. 2. HSC scATAC-seq training curves

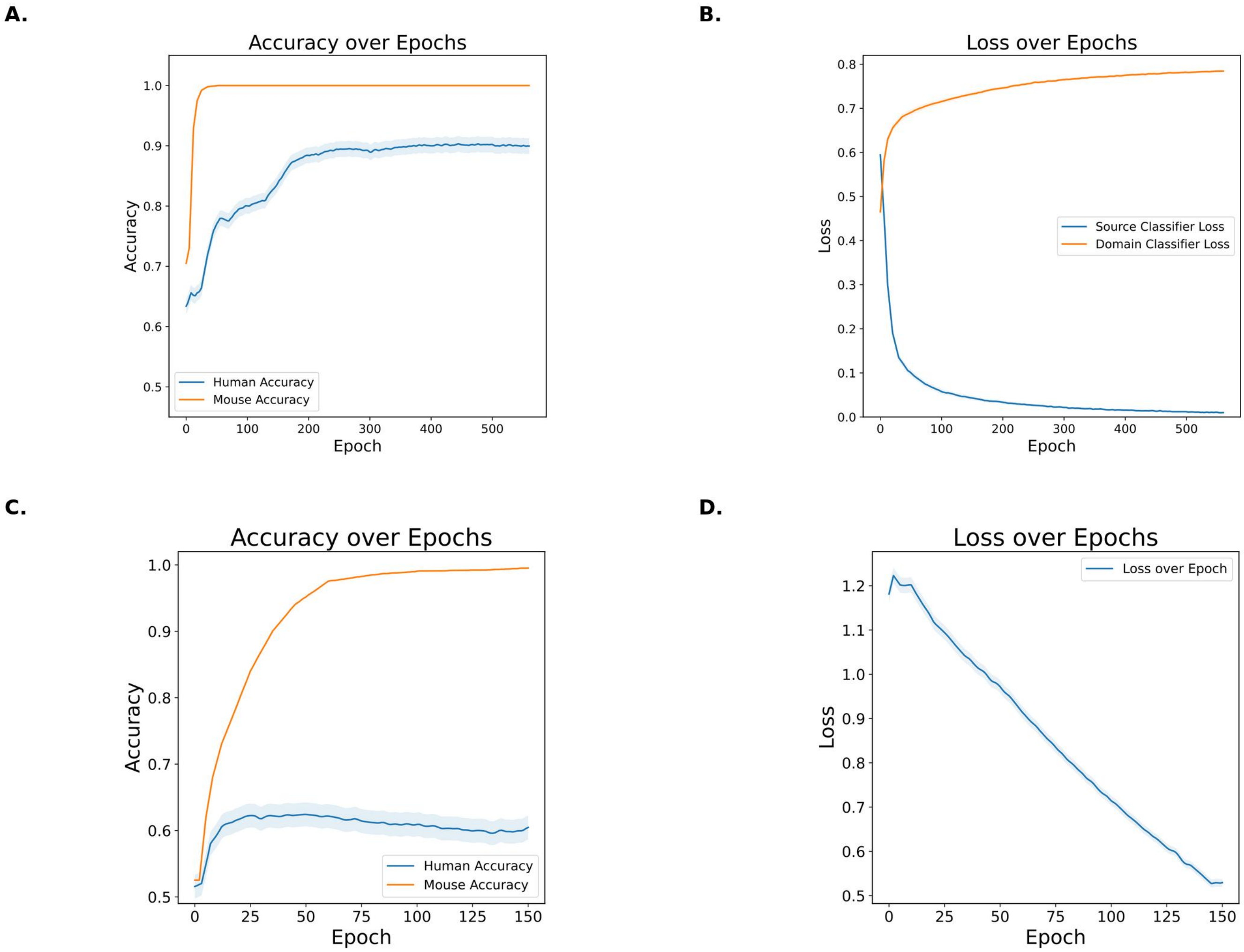

### Extended Data Fig. 3. CD8+ T-cell scRNA-seq training curves

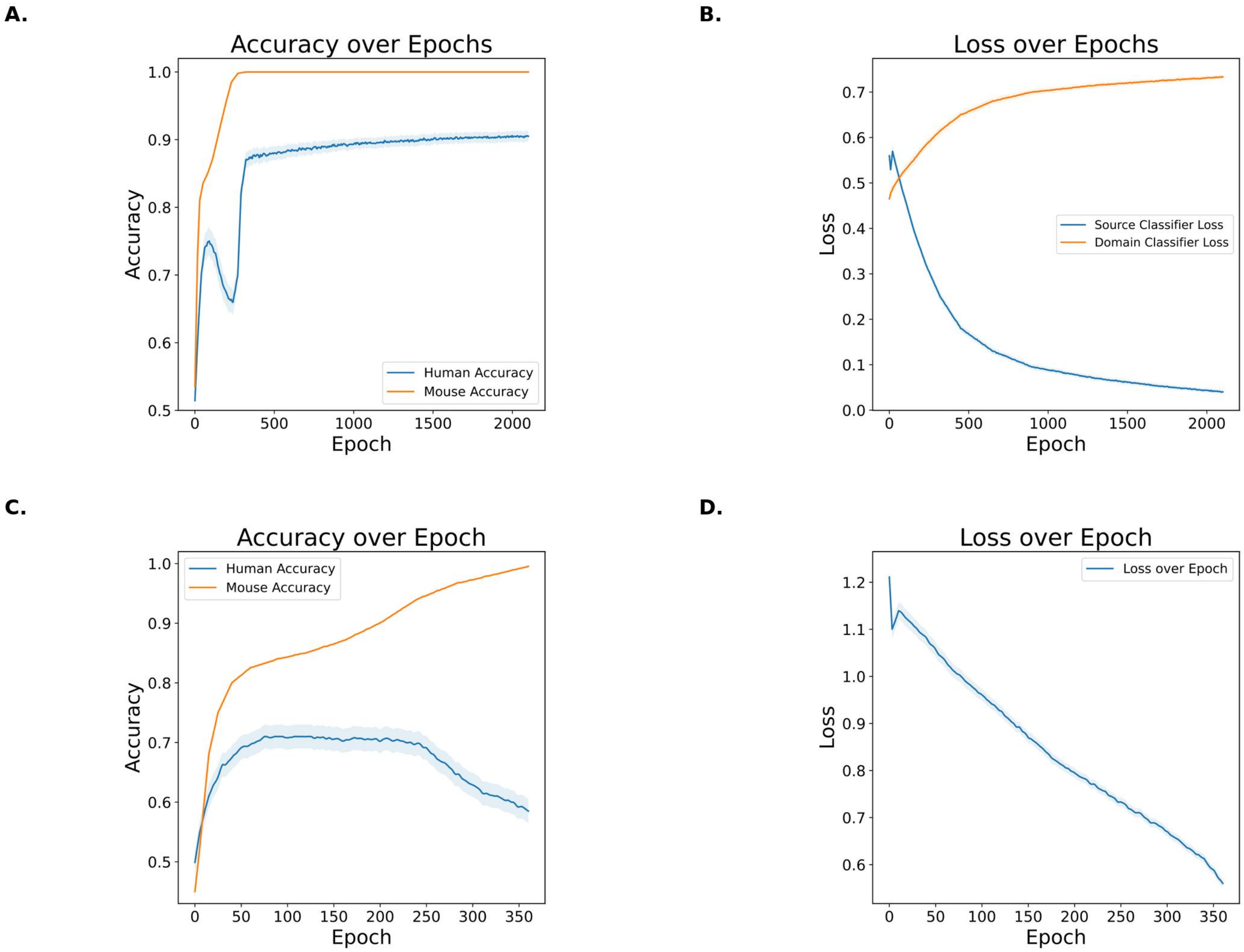

### Extended Data Fig. 4. HSC scATAC-seq DANN embedding evolution

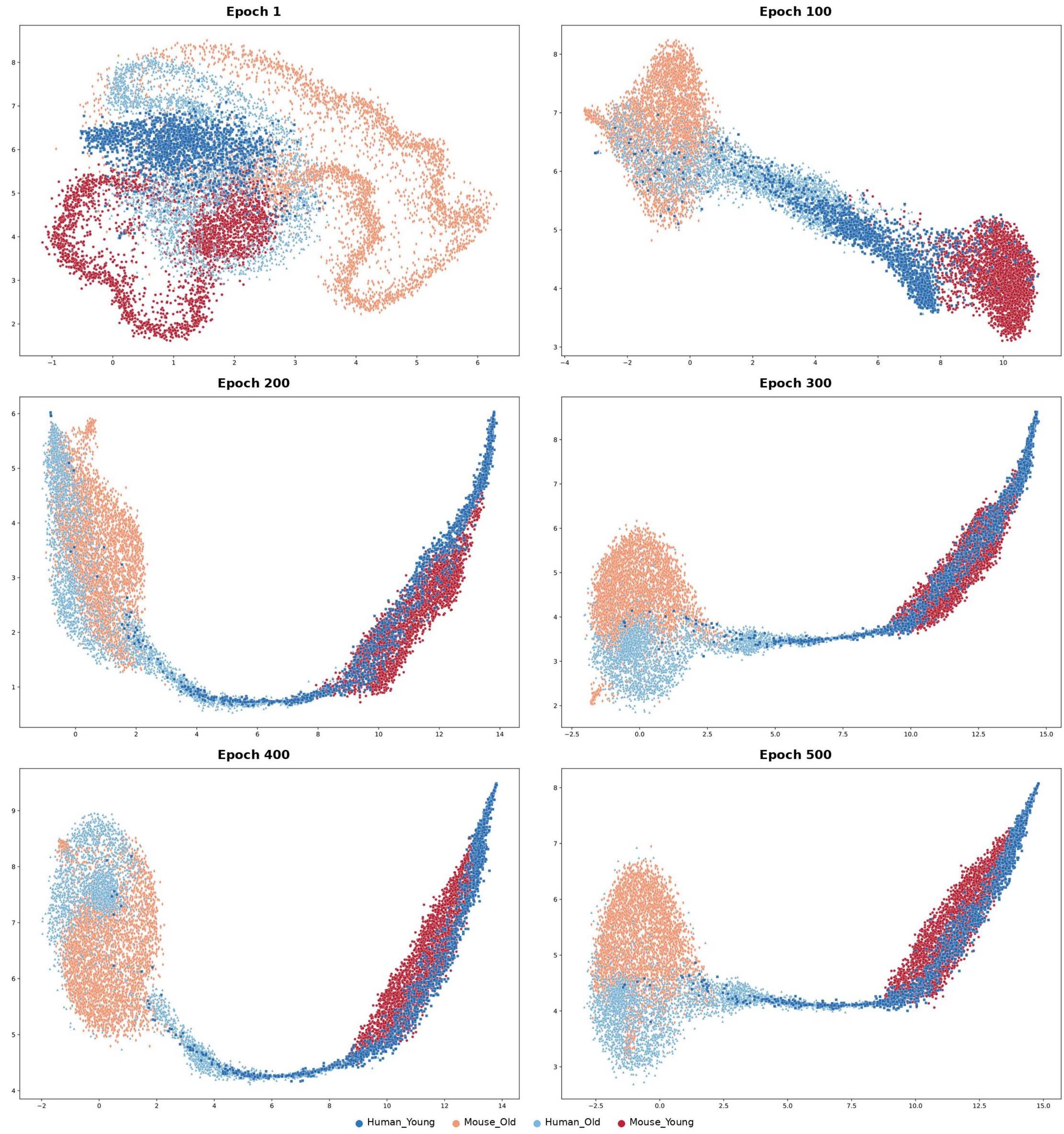

### Extended Data Fig. 5. HSC scRNA-seq DANN embedding evolution

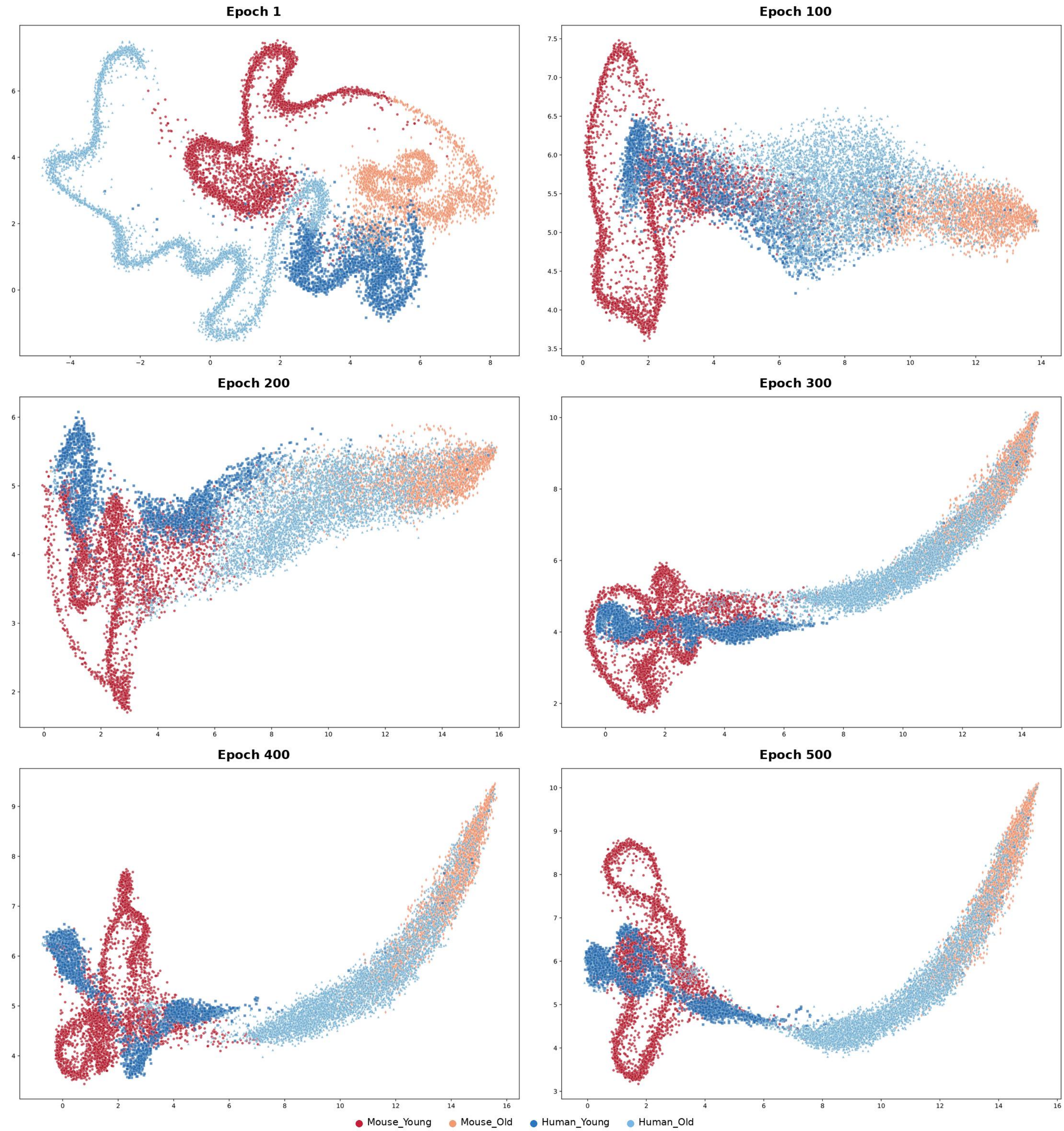

### Extended Data Fig. 6. CD8+ T-cell scRNA-seq DANN embedding evolution

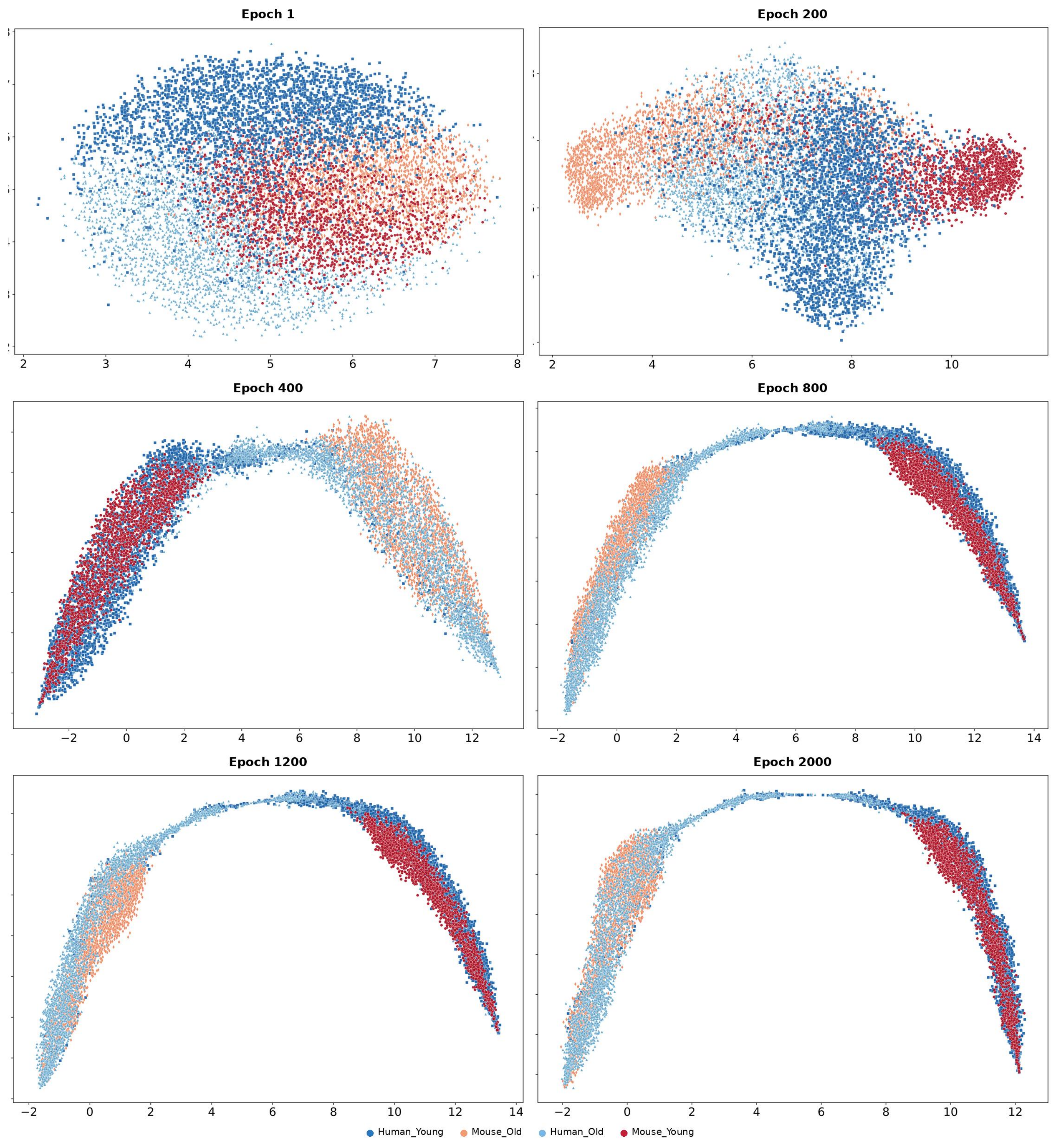

### Extended Data Fig. 7. Attribution reproducibility and consensus analysis

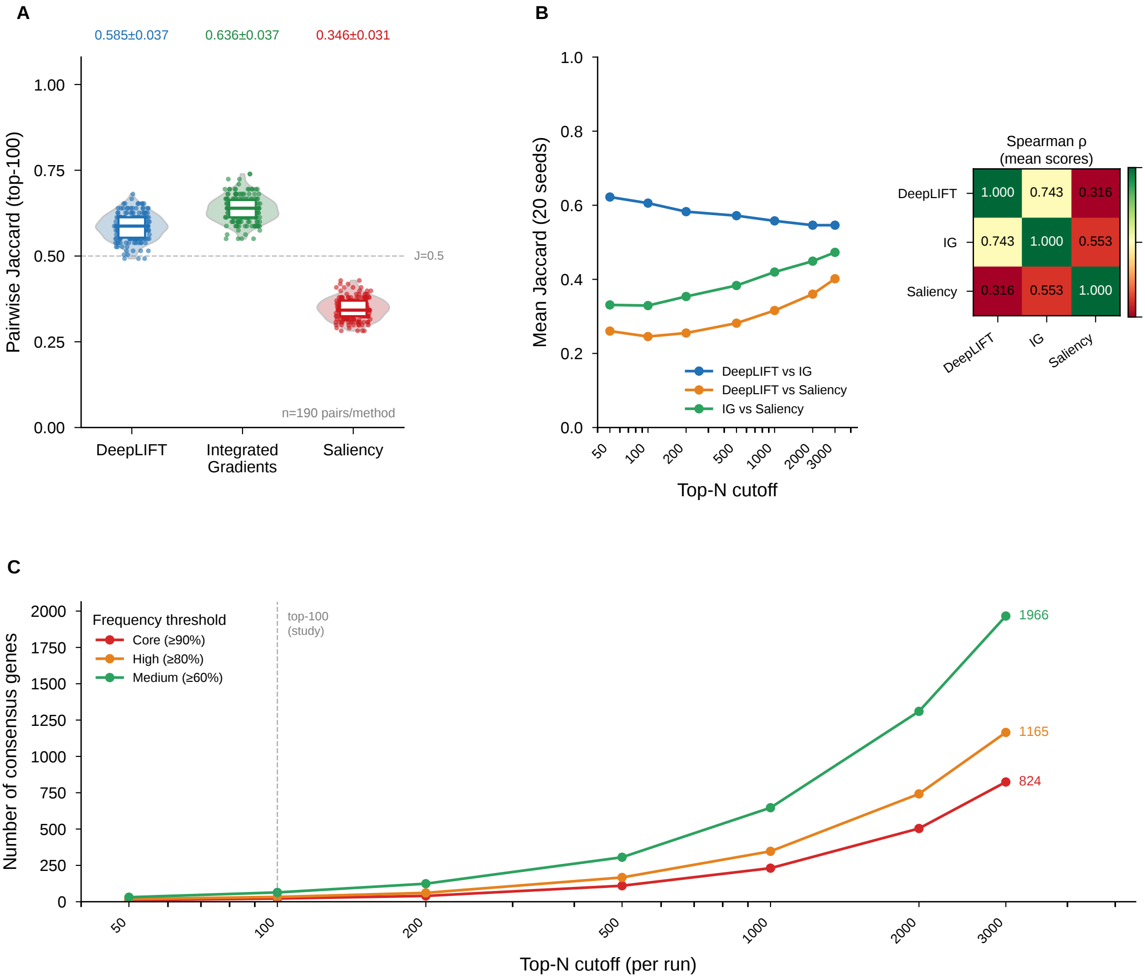

### Extended Data Fig. 8. Model ablation and architecture sensitivity

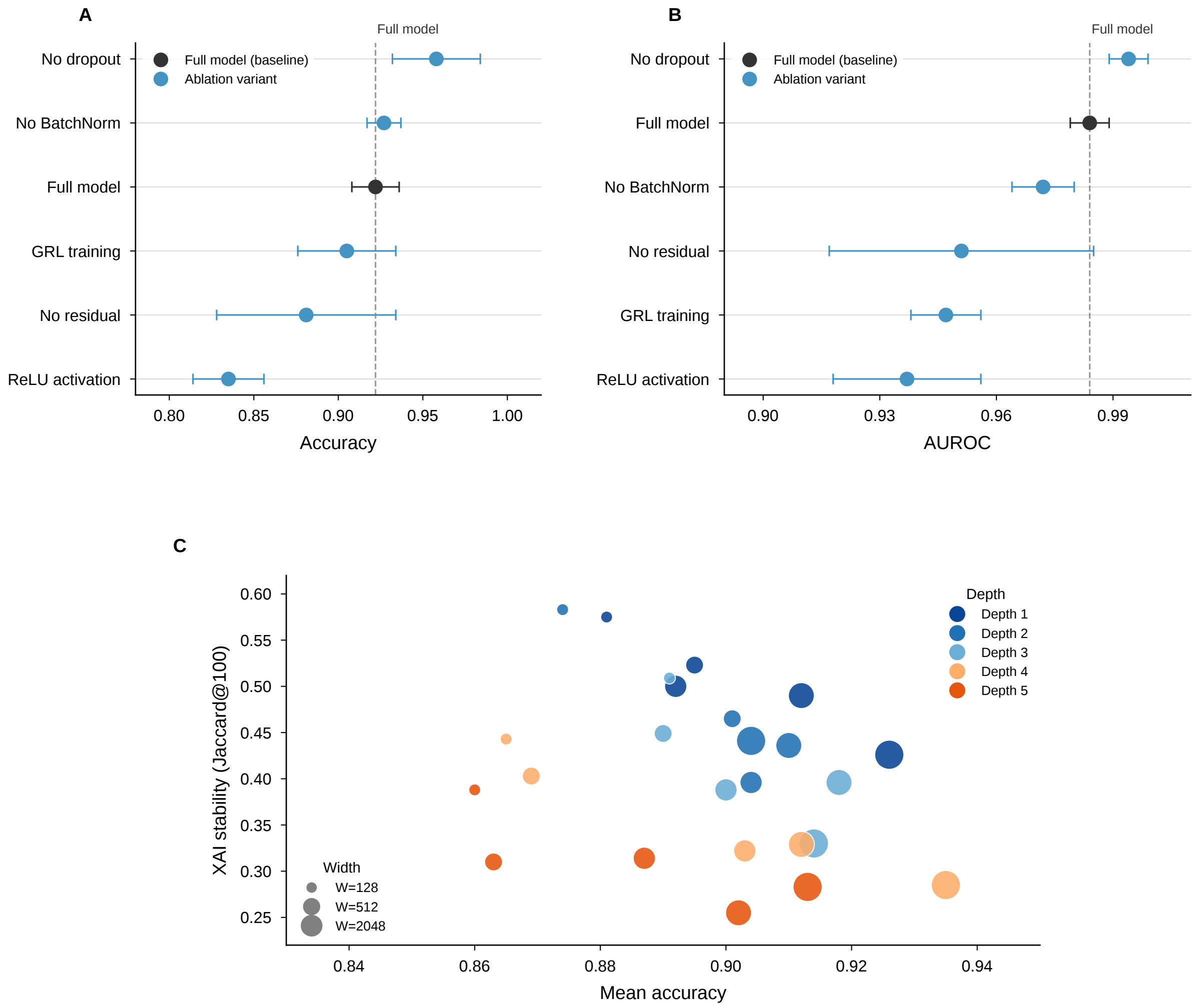

### Extended Data Fig. 9. CD8+ T-cell cross-species attribution analysis

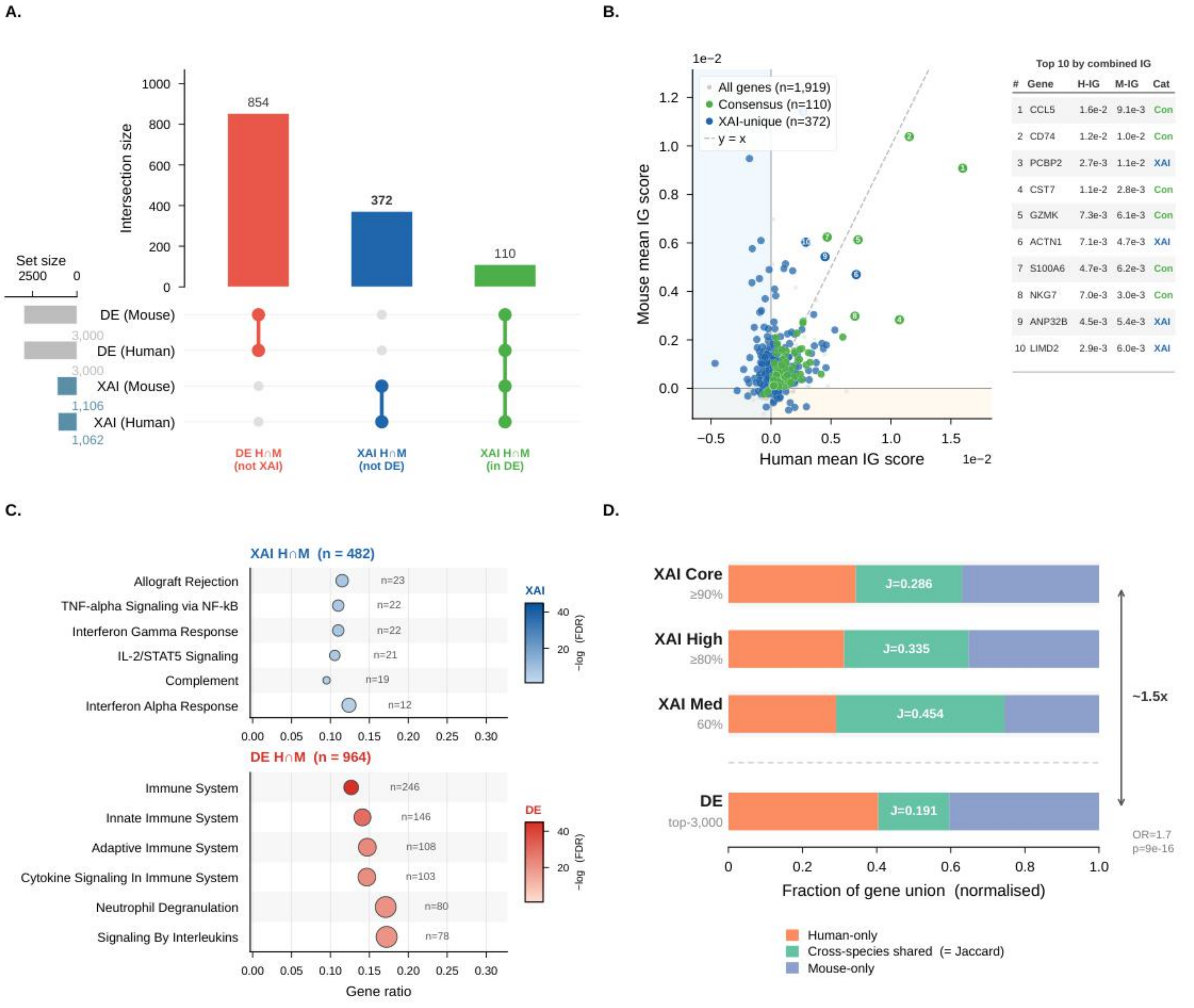
